# Developing a predictive model for an emerging epidemic on cassava in sub-Saharan Africa

**DOI:** 10.1101/2022.05.13.491768

**Authors:** David Godding, Richard O. J. H. Stutt, Titus Alicai, Phillip Abidrabo, Geoffrey Okao-Okuja, Christopher A. Gilligan

## Abstract

The agricultural productivity of smallholder farmers in sub-Saharan Africa (SSA) is severely constrained by pests and pathogens, impacting economic stability and food security. Since 2004, an epidemic of cassava brown streak disease (CBSD) has been spreading rapidly from Uganda, with the disease causing necrosis of the edible root tissue. Based on sparse surveillance data, the epidemic front is currently believed to be at least as far west as central DRC and as far south as Zambia. The DRC is the world’s highest per capita consumer of cassava and future spread threatens production in West Africa which includes Nigeria, the world’s largest producer of cassava. Here, we take a unique Ugandan CBSD surveillance dataset spanning 2004 to 2017 and develop, parameterise, and validate a landscape-scale, spatiotemporal epidemic model of CBSD at a 1 km^2^ resolution. While this paper focuses on Uganda, the model is designed to be readily extended to make predictions beyond Uganda for all 32 major cassava producing countries of SSA, laying the foundations for a tool capable of informing strategic policy decisions at a national and regional scale.

## 1 Introduction

A principal challenge in dealing with emerging epidemics and pest infestations of agricultural crops is to estimate the current extent of infection and the rates of spread across heterogeneous landscapes. The challenges are particularly acute for epidemics that impact smallholder agriculture in sub-Saharan Africa (SSA), where many staple crops are under threat from emerging pests and pathogens. Examples include maize lethal necrosis [1], fall armyworm [2], banana bunchy top disease [3], wheat rusts [4, 5], cassava viruses [6], and desert locust [7].

Cassava is the second most important source of calories in SSA after maize [8]. Cassava brown streak disease (CBSD) is caused by cassava brown streak ipomoviruses (CBSIs); ssRNA *Ipomoviruses* of the family *Potyviridae* that are epidemiologically equivalent. The disease poses one of the most significant threats to cassava production in SSA, causing necrosis of the edible root tissue [9, 10]. The CBSIs are spread by an insect vector, the whitefly *Bemisia tabaci* [11], with additional spread by trade movement of virus-infected cuttings used for planting [12].

Following an initial report of CBSD in Uganda in 2004 [13], the disease rapidly spread throughout Uganda [14] to surrounding countries, including Rwanda [15], Burundi [16], western Kenya [17], lake-zone Tanzania [18], eastern DRC [19], and Zambia [20]. The disease has more recently been reported as far west as the north-central province of Tshopo, DRC [21]. Continued westward spread and the risk of direct introduction via the movement of planting material poses a major threat to food security and economic stability in Central and West African countries.

An initial step in predicting the onward spread of the pathogen is to estimate transmission and dispersal parameters at landscape scales. We do this by fitting and validating a stochastic, spatially-explicit metapopulation epidemic model of CBSD spread in Uganda, at a 1 km^2^ resolution, using a unique multi-year country-wide surveillance dataset [14] that documents the progressive spread of CBSD throughout Uganda from a few isolated initial infected sites near Kampala. The model takes account of the spatial distribution and connectedness of the cassava crop and variability in the abundance of the insect vector throughout Uganda. Estimating parameters from sparse spatiotemporal data is extremely challenging and an area of active research. We use approximate Bayesian computation (ABC), which does not require the explicit definition of the likelihood and is well adapted to dealing with unobserved data [22] when inferring sequences of infection spread across a heterogeneous landscape. However, a central challenge is specifying summary statistics that capture as much information as possible about the dynamics of the system in the simplest possible form [22–24].

Specifically, we address the following questions considering epidemic spread of CBSD:

- Can a simple epidemiological model structure capture the fundamental dynamics of the CBSD epidemic?
- Can transmission rates and dispersal parameters be estimated from disjoint snapshots of annual surveillance data?
- Can the parameterised model predict the future spread of the virus in successive years within Uganda?

## 2 Results

### 2.1 Incorporating data-driven model layers

A spatially explicit, stochastic SI metapopulation epidemic model provided the framework for the spread of infection, and by implication disease, within and between rasterised cells in the landscape. A data-driven host landscape layer was generated to account for the spatial heterogeneity of cassava production. The host landscape layer was derived by converting the CassavaMap model [25] from production volume in tonnes per km^2^ to the number of fields per km^2^ (Figure 1a).

**Figure 1:**
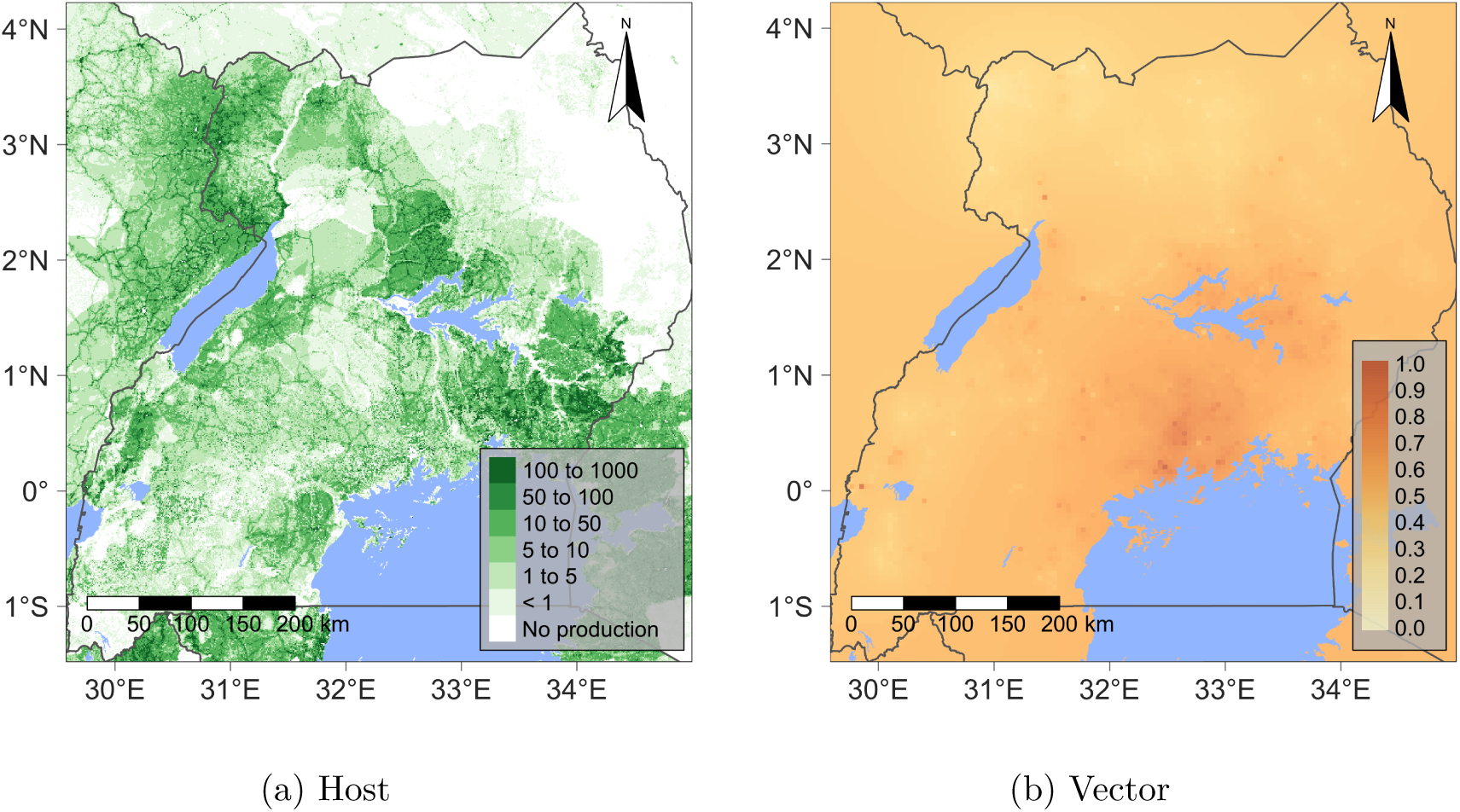
Maps representing the rasterised data-driven model layers: (a) The model host landscape, representing the number of cassava fields at a 1 km^2^ resolution derived from CassavaMap [25]. (b) The vector abundance layer represents the relative abundance of *B. tabaci* at a 5 km resolution derived from the Ugandan CBSD field surveys [14].

Through an iterative process of model development (Supplementary Methods S1.1), the model was extended to incorporate an additional epidemiologically important data-driven spatial layer accounting for the variation in the abundance of the vector, *B. tabaci*, across the Ugandan landscape (Figure 1b). The rasterised vector abundance layer was generated from the *B. tabaci* count data collected as part of cassava field surveys [14].

### 2.2 Parameterising the spatiotemporal epidemic dynamics

The Ugandan surveillance data, *d*_*real*_, were divided into two distinct datasets. The training dataset, 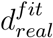, consisting of data from 2004 to 2010 inclusive, and validation dataset,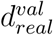, consisting of the remaining data from 2011 to 2017. We applied ABC rejection to estimate three model parameters using 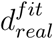 a dispersal kernel exponent, *α*, a transmission rate, *β*, and the proportion of dispersed inoculum that remains in the source cell, *p*. The posterior probability distribution for these parameters was calculated as the number of simulations generating simulated surveillance data, 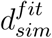, sufficiently close to the real-world surveillance data, 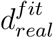, normalised relative to the sampling density (Supplementary Methods S1.2) [24, 26].

In order to quantify the distance between the real-world training data, 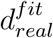, and the simulated surveillance data covering the same time period, 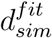, we constructed three epidemiologically-informed summary statistics (see Parameter estimation in Methods). The three summary statistics capture different aspects of the spatiotemporal characteristics of the epidemic: *S*_*nat*_ and *S*_*cen*_ are based on calculations of the proportion of survey points in different regions that are reported as positive each year. A third statistic, *S*_*grid*_, explicitly tracks the spatial expansion of the epidemic throughout Uganda in terms of the year of first CBSD detection in each cell of a regular grid covering the full extent of Uganda.

The statistic, *S*_*cen*_, is designed to capture the local bulk-up dynamics of the epidemic in a small, densely sampled area in central Uganda surrounding Kampala, with dimensions: *x*_*min*_ = 32.20, *x*_*max*_ = 33.21, *y*_*min*_ = 0.09, *y*_*max*_ = 1.16 in the WGS 84 coordinate system. The statistic, *S*_*nat*_, covers the remaining non-overlapping spatial extent of Uganda and captures the regional bulk up rate (Figure 2a). For convenience, we refer to the combination of summary statistics, *S*_*cen*_, *S*_*nat*_ and tolerances, *ϵ*_*cen*_, *ϵ*_*nat*_ as *S*_*inf*_ and *ϵ*_*inf*_ respectively.

**Figure 2:**
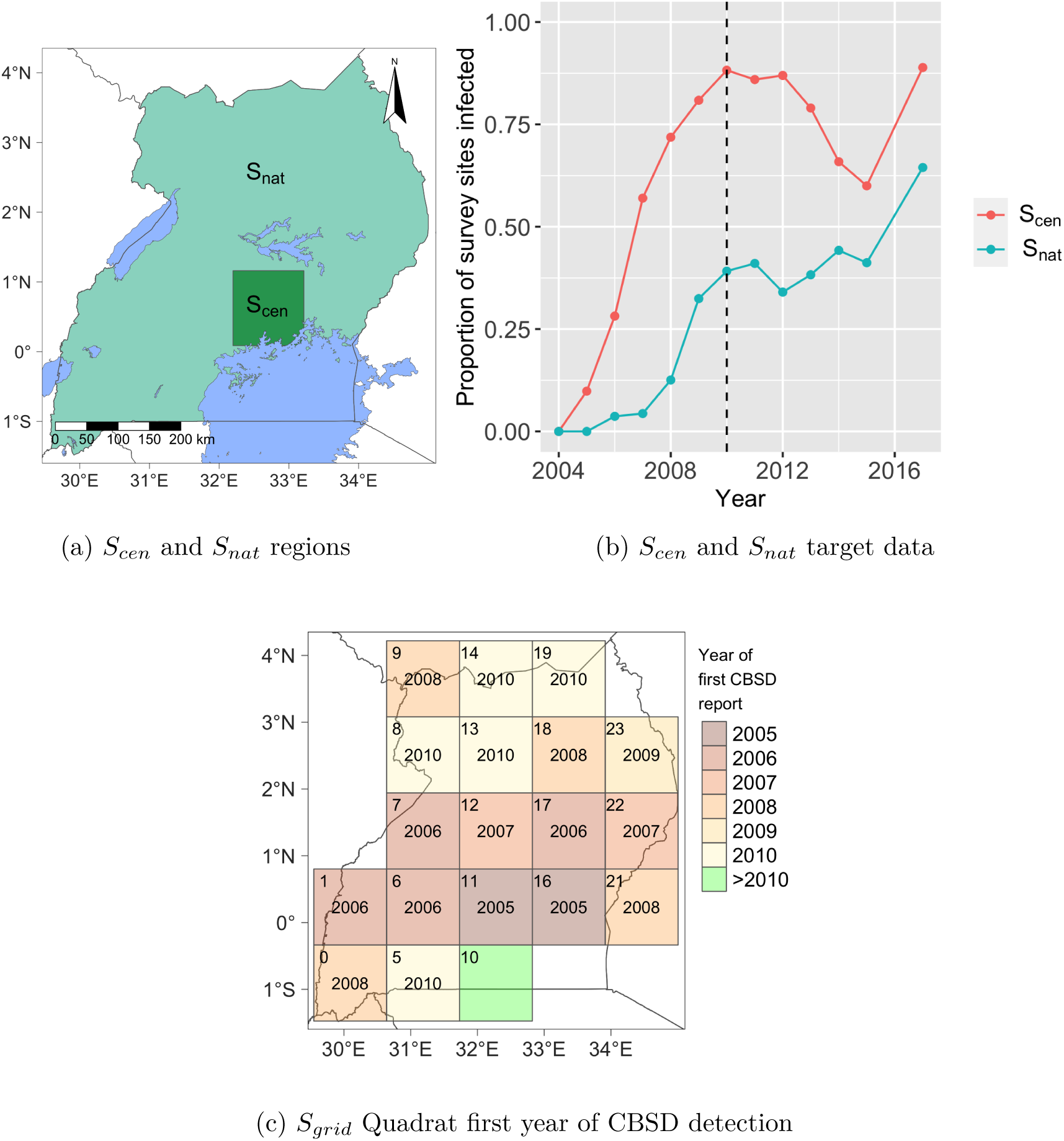
Overview of the three summary statistics used for ABC parameter estimation. Both *S*_*cen*_ and *S*_*nat*_ statistics were derived by calculating the proportion of survey points in a given year that were reported as positive within a given region: (a) represents the two non-overlapping areas of Uganda covered by *S*_*cen*_ and *S*_*nat*_ and (b) summarises the values derived when applying *S*_*cen*_ and *S*_*nat*_ to the Ugandan national survey data covering the period 2004 to 2017. The dotted black line indicates the divide between the fitting data, 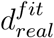, and the validation data, 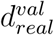 (c) Overview of the summary statistic *S*_*grid*_ highlighting the survey year, up to and including 2010, in which a CBSD infected field was first detected in a given quadrat. If no positive surveys were reported prior to 2010, as in the case of quadrat 10, the quadrat is shaded green. If no surveys were carried out prior to 2010, the quadrats have been excluded from the plot. Quadrat indices are shown in the top left corner of each quadrat.

The third statistic, *S*_*grid*_, is derived by dividing the latitude/longitude extent of Uganda into a 5×5 grid of quadrats. For a given simulation, the statistic is scored as the proportion of the quadrats where infected fields are detected in either the same year as the real-world surveillance data or *±*1 year either side, or in the case where all surveys in a given quadrat were negative for CBSD, the simulation must remain negative in all simulated surveys (Figure 2c).

The ability of the summary statistics to recover known parameter values was first tested using synthetic data for the spread of CBSD across the Ugandan landscape. Preliminary analyses also showed that the statistics were best used in combination, and guided the selection of appropriate tolerances for each statistic (Supplementary Methods S1.3)

The posterior distribution is derived from 1440 simulations that passed the fitting criteria (i.e. tolerances applied to the three summary statistics) from a total of 233,600 fitting simulations (Figure 3). For the kernel scale parameter, *α*, and log of transmission rate, ln(*β*), the posterior distribution covers a clear and distinct region of highest posterior density, with a correlation between shorter dispersal distances requiring higher transmission rates and vice versa. Within the credible ranges of *α* and ln(*β*), the third parameter, *p*, governing the amount of inoculum that remains in the source cell, lies almost exclusively below 0.8 and is concentrated around 0.12.

**Figure 3:**
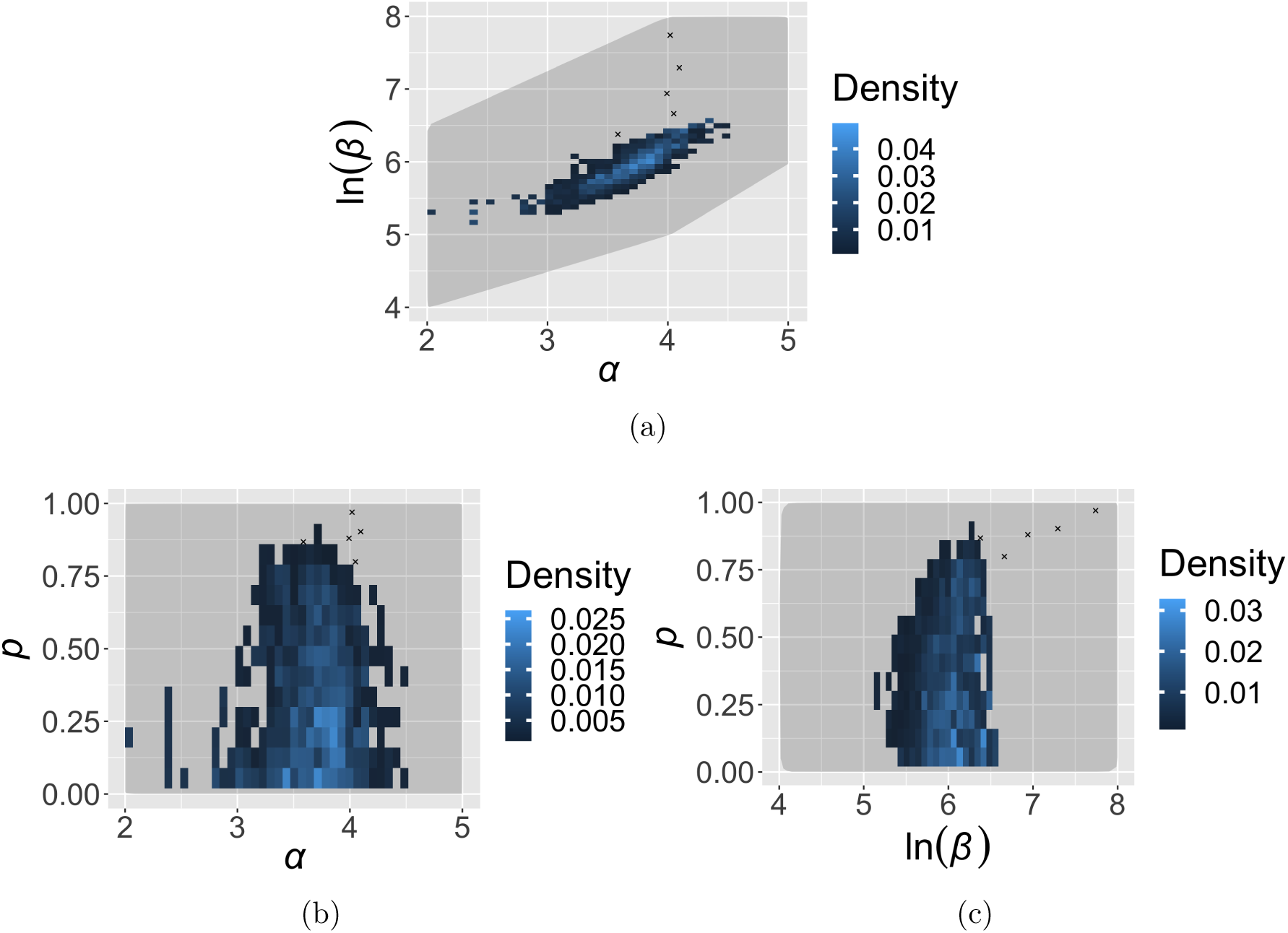
Posterior distribution of the three parameters estimated using the fitting data from 2004 to 2010, 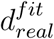. The parameters are the transmission rate, *β*, the kernel exponent, *α*, and the proportion of dispersed inoculum that remains in the source cell, *p*. The posterior is composed of 1440 fitting simulations that met the fitting criteria out of a trial of 233,600 fitting simulations. Five outliers, indicated by black crosses, were excluded from sparsely sampled parameter space (Supplementary Figure 7).

### 2.3 Simulating the impact of a disease management programme

The summary statistic *S*_*cen*_ highlights a clear but temporary reduction in the proportion of field surveys reporting CBSD from 2013 to 2015 in the small central area surrounding Kampala (Figure 2b). The reduction in the intensity of the epidemic in this region was the result of a number of projects that disseminated a total of 40 million virus-free cassava cuttings, focusing on four Ugandan districts: Luwero, Mukono, Nakasongola, Wakiso (Figure 4a), with surveillance in each district capturing the same characteristic pattern of temporary decline (Figure 4b) [27, 28]. Beyond this high level information, specific details on precise location and timing of the different programmes that disseminated clean planting material are not available.

**Figure 4:**
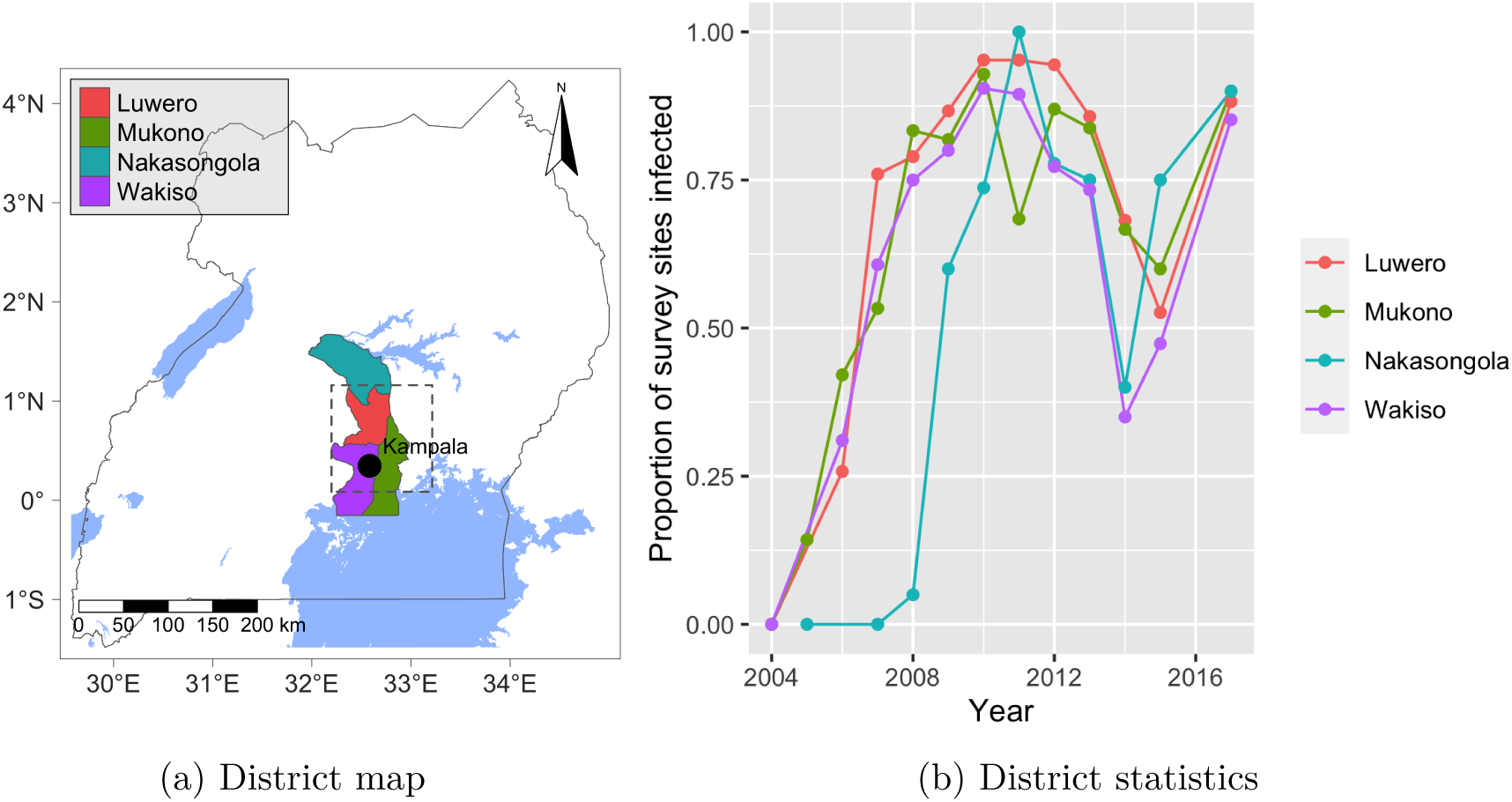
Overview of the districts where clean seed dissemination programmes were primarily carried out: (a) shows the location of the districts in Uganda as well as the region covered by *S*_*cen*_ (dotted line) and (b) summarises the proportion of field surveys that reported CBSD in a given year from each district.

We implemented a process equivalent to the clean seed programmes in the model via three discrete rounds of *I → S* replacement at the start of the 2013, 2014, and 2015 growing seasons. The parameter, *r*_*clean*_, defines the proportion of CBSI infected cassava fields across the four districts that should be replaced by virus-free planting material in each of the three rounds, with the exact fields being selected at random. A value for *r*_*clean*_ of 0.15 was selected based on a parameter sweep to identify the value that best fitted the observed impact of the clean seed programmes in *S*_*cen*_ (Supplementary Methods S1.1.4). The clear correspondence of the simulated clean seed programme to the real-world observations, viewed through the lens of the summary statistic, *S*_*cen*_, is illustrated during model validation.

### 2.4 Validating predictions of epidemic spread

An ensemble of 10,000 simulations were run from 2004-2010 by sampling from the posterior parameter distribution to generate initial conditions for the validation simulations. Of these 10,000 simulations, 65 met the fitting criteria, which were then used as initial conditions for the system state at the start of 2011. These simulations were then run for the validation period of 2011-2017 and their correspondence to the validation data, 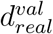, was assessed by applying the validation criteria (i.e. tolerances applied to the summary statistics, *S*_*inf*_, during the validation period). All 65 validation simulations passed the validation criteria, resulting in a validation score of 100% (Figure 5).

**Figure 5:**
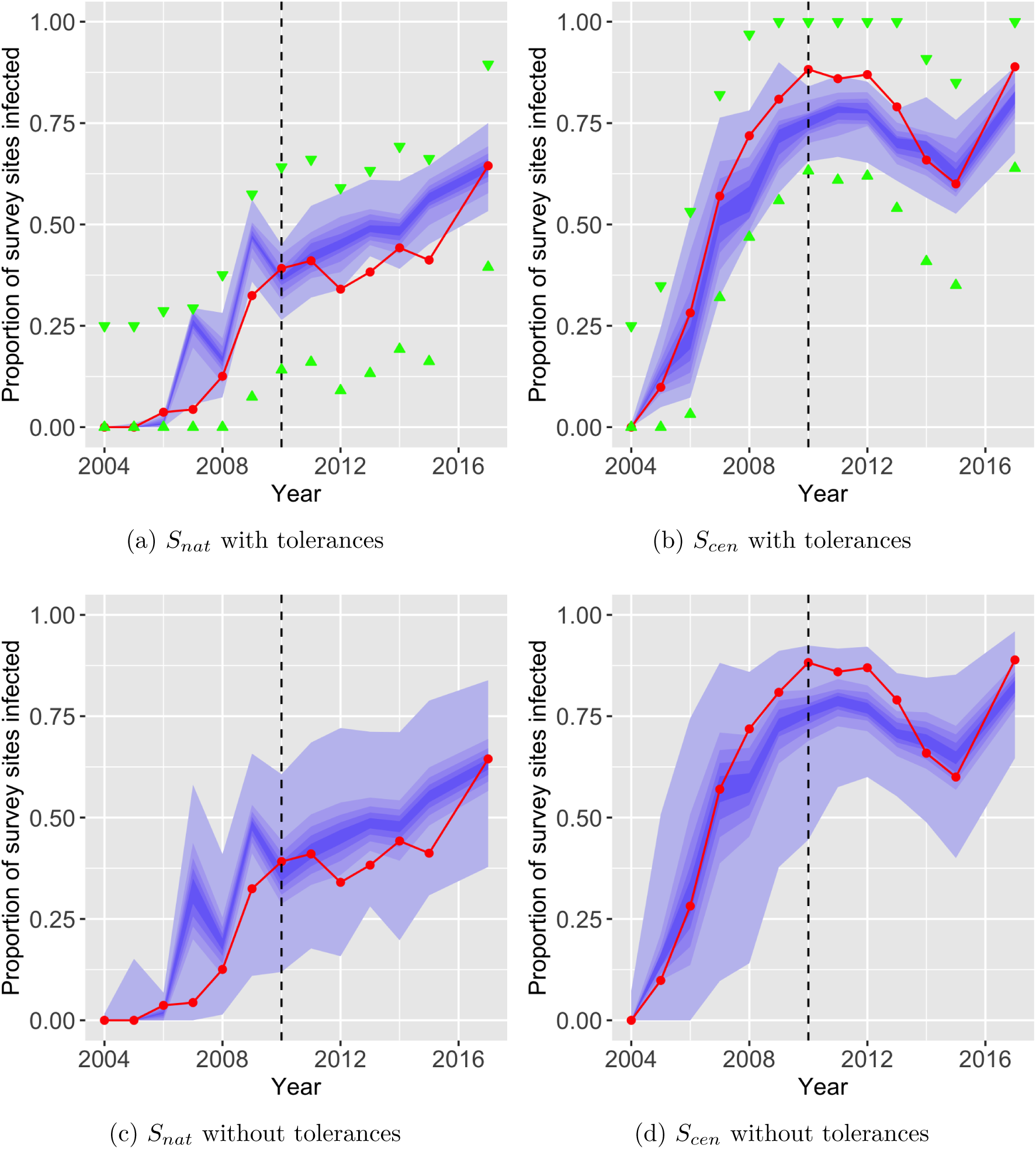
Time series probability distributions for the statistics (a) *S*_*nat*_ and (b) *S*_*cen*_ for the subset of validation simulations that pass within the tolerances of the fitting and validation criteria. (c) and (d) show the same statistics but without applying any tolerances to illustrate the unconstrained behaviour of the parameterised model. The red line indicates the target value of each statistic derived from surveillance data. Tolerances are indicated by green arrows. The central blue band is the median *±*10% and each gradation beyond is a further *±*10% from the median. The dotted black line indicates divide between 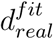 and 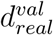.

Figure 6 illustrates the spatial structure of the simulated survey data from a single validation simulation, illustrating the strong yearly correspondence between simulated and real-world surveillance from 2005 to 2017 in terms of the spatial distribution of fields reported as present/absent for CBSD and the local density of CBSD positive surveys.

**Figure 6:**
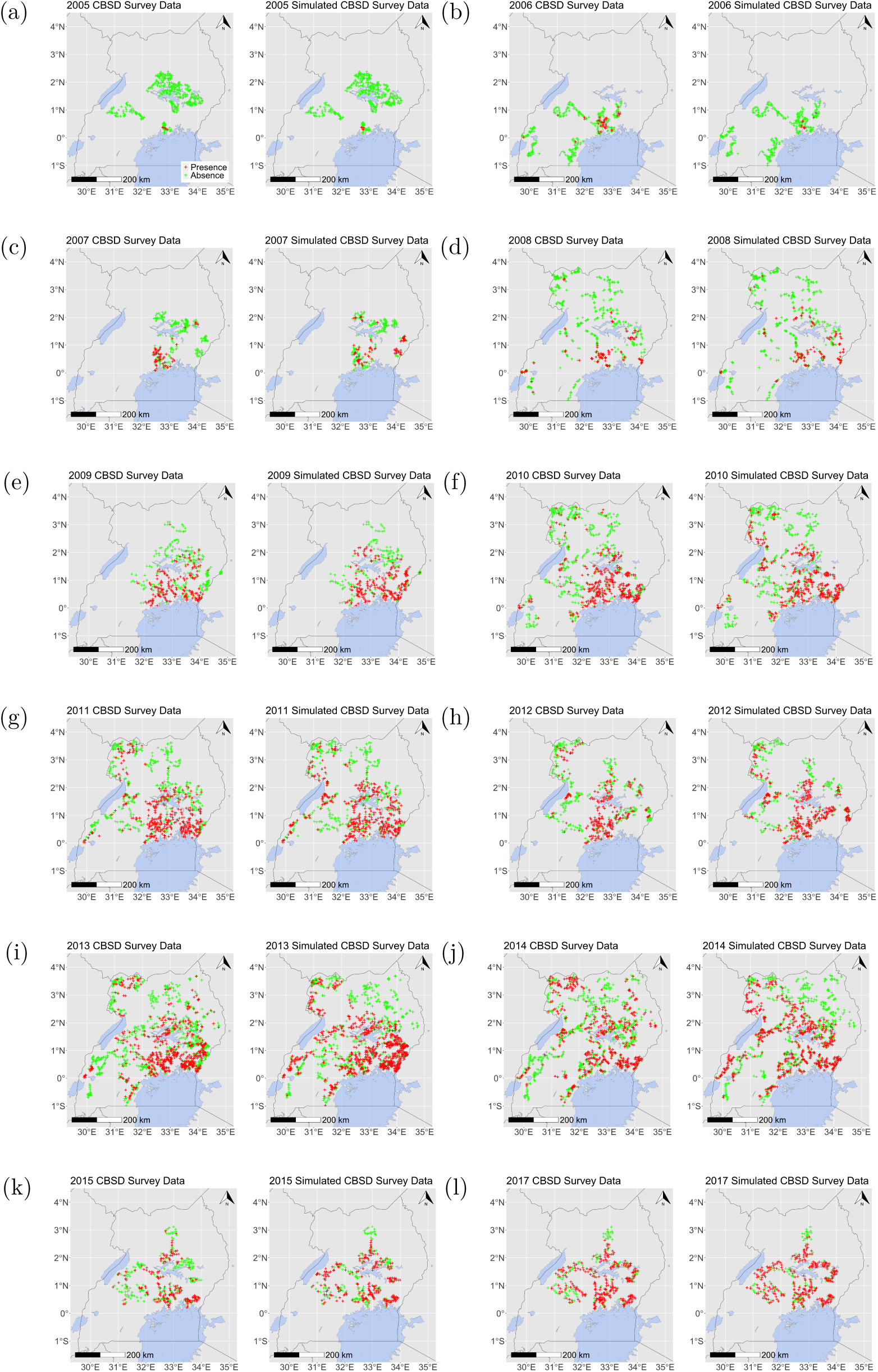
Comparison of a single validation simulation that passes the fitting and validation criteria with the real-world surveillance data from 2005-2017. We define the fitting criteria as the values selected for the three tolerances for parameter estimation using Ugandan survey data from 2004-2010: 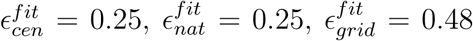 and the validation criteria as the comparison of the simulated surveillance data to the Ugandan survey data covering the validation period from 2011-2017, 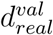, using the two *S*_*inf*_ statistics with tolerances 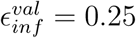 = 0.25. Red crosses indicate an observation of CBSD at the field-level. Green crosses indicate no CBSD observed.

## 3 Discussion

It is essential that policy makers across the cassava producing countries of sub-Saharan Africa have a clear understanding of the current state of the CBSD epidemic and the likely future spread to assist in deciding when and how to mitigate the impact of the CBSD epidemic and minimise future spread. However, surveillance data for the ongoing CBSD epidemic are extremely sparse, especially further west of the now endemic regions in East Africa due to complex geographic, political and financial constraints. Moreover, until now, no landscape-scale spatial epidemic models of CBSD existed to extrapolate beyond surveillance data and provide a shared quantitative framework to assist policy formulation. To this end, we have presented the development, parameterisation and validation of a landscape-scale stochastic model of the CBSD epidemic in Uganda. The fitted model shows strong correspondence to the validation dataset, 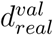 (Figure 5 and 6). Importantly, whilst this study focusses on Uganda, the data-driven host and vector input layers are readily extended to the entirety of sub-Saharan Africa. Moreover, the success of the model in simulating the impact of the localised deployment of virus-free planting material in the region around Kampala between 2013 and 2015 provides initial evidence for the flexibility of the model to predict and analyse the impacts of management scenarios (Figure 5).

The available data on the CBSD pathosystem are both spatially and temporally sparse. The pathogen has two dispersal mechanisms: vector-borne spread and human-mediated movement of infected planting material. In the absence of targeted data collection to distinguish between the two mechanisms, we have parameterised a single dispersal kernel that represents the net effect of both underlying forms of dispersal across the Ugandan cassava landscape. The parameterised model proved sufficient to characterise the spread of the pathogen in Uganda, with a marked correspondence between simulated and real-world surveillance data for both the data fitting period, 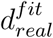(2004-10), and validation period, 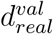(2011-2017) (Figure 6). We recommend, however, that future work should focus on disentangling the two dispersal mechanisms.

Due to an absence of cheap, reliable in-field diagnostics for CBSD, the survey protocol was based on above ground foliar symptoms, which likely leads to an underestimation of the true prevalence at the plant level [29]. The impact of this underestimation is mitigated by modelling spread at the field level with data be-ing aggregated across the 30 plant sample to a field level presence/absence record. Moreover, we allow for false negative biases in the model structure to account for the limitations of the protocol, such as the limited number of plants surveyed per field and biases in cultivar selection (Supplementary Methods S1.1.1).

The model is modulated by data for the spatial distribution of the host crop and the insect vector across the Ugandan landscape, thereby improving model fit (Supplementary Methods S1.1.3). The host landscape layer provides the best available model of cassava production at a 1 km2 resolution across sub-Saharan Africa [25], providing a single temporal snapshot of the amount of cassava being grown. We therefore assume a constant distribution of cassava production across Uganda over time. We also assume that all cassava is equally susceptible to infection by CBSIs, which is consistent with currently available evidence [30, 31]. Similarly, evidence to date does not indicate major differences in either the yield impact or geographic distribution of the two CBSIs [32]. Hence, for the purposes of this study, we did not distinguish between the two.

The vector abundance layer is derived from the interpolation of values from the Ugandan cassava field surveys that quantified *B. tabaci* abundance. It is important to note that the uncertainty in vector abundance layer values is higher in regions with lower spatiotemporal surveillance density. As with the host landscape, the vector abundance layer provides a single atemporal snapshot, therefore assuming the vector abundance remains stable over time and the different species in the *B. tabaci* complex are equivalently capable of transmitting CBSIs. There is an ongoing debate over the extent to which there is an interaction between the cassava epidemics of CBSD and cassava mosaic disease (CMD) and changes in local vector abundance or the specific abundance of species within the *B. tabaci* complex [33, 34]. Extensive experimental and modelling work would be necessary to improve our understanding of the significance of the local composition and abundance of the *B. tabaci* species complex on the spread of the epidemic. Despite this complexity, it is clear that the incorporation of the vector abundance layer improved the predictive power of the model (Supplementary Methods S1.1).

The model presented in this study represents a significant advancement in our ability to predict the spread of the CBSD epidemic and simulate disease management scenarios. Importantly, the model has the potential to act as an overarching quantitative framework to assist in addressing a number of vital questions: how can CBSD endemic countries minimise the impact and reduce the prevalence of CBSD; when will CBSIs spread to currently unaffected countries in West Africa; and how can these not yet affected countries optimise surveillance for early detection and prepare to control an outbreak.

## 4 Methods

We developed, parameterised and validated a stochastic metapopulation epidemic model for CBSD in Uganda at a 1 km^2^ resolution via an iterative process of model development (Supplementary Methods S1.1). The model integrates a host landscape of cassava production and the relative spatial abundance of the insect vector, *B. tabaci* (Figure 1). We simulated the spread of CBSD across the host landscape as a spatially explicit, SI (Susceptible-Infected) epidemic via a discrete event, continuous time stochastic process using an optimised Gillespie algorithm [35, 36]. An SI compartmental structure was selected as cassava is a vegetatively propagated crop, so infection persists from one harvest to the next planting [37]. The model was parameterised and validated using annual surveillance data for the spread of CBSD in Uganda [14] (Supplementary Figure 1).

### 4.1 CBSD surveillance data

Surveillance of the CBSD epidemic in Uganda was carried out in annual field surveys since the start of the epidemic in 2004 through to 2017, with the exception of 2016 [14]. In a given year, surveyors visited between 253 and 1250 fields, with a mean of 587. The spatial distribution of surveys was not uniform. In some years, surveys were carried out in relatively small regions, whereas in other years surveys were more evenly distributed throughout the country. The same fields were not revisited across multiple years. The survey protocol for a given field involved surveyors randomly selecting 30 plants of the dominant cultivar across two diagonal transects and recording the severity or absence of visual CBSD foliar symptoms, along with the number of individual *B. tabaci* on the upper five leaves.

From the perspective of assessing disease presence at the field level, two factors likely resulted in a degree of systematic under-reporting of disease. Firstly by sampling only the dominant cultivar surveyors did not record disease on non-dominant varieties. Secondly, a sample size of 30 plants is small relative to a total of approx-imately 1000 plants in a field of 0.1 ha. However, the dataset contained additional information for a subset of fields that allowed us to estimate a false negative reporting rate. For data collected from 2009 to 2014, surveyors reported whether CBSD disease symptoms were observed anywhere in the field, as opposed to just on the dominant variety or the 30 plant sample. Based on an analysis of these records, the average false negative under-estimation rate was estimated to be approximately 0.15 (Supplementary Methods S1.1.1).

### 4.2 Model structure

For the host lanscape layer, we assume an average per field cassava yield of 10 tonnes per hectare [38] and an average field size of 0.1 ha [39–41]. The CassavaMap model used two forms of input data: human population data and regional cassava production statistics. For each region, the total production volume was allocated in proportion to the number of inhabitants per km^2^ with the exception of spatial locations with populations greater than 5000 inhabitants per km^2^, which were excluded to avoid the allocation of production to urban areas. The model caps production at 1000 tonnes per km^2^ [25].

For the vector abundance layer, field-level vector abundance mean values were collapsed across all survey years to create a single atemporal dataset. Field-level means were then capped to a maximum credible mean value of 100. Inverse distance weighted (IDW) interpolation was applied with a power value of 1.0, generating a rasterised layer with a 5 km resolution. A linear relationship between *B. tabaci* count and field-level infectiousness and susceptibility was selected, reaching saturation at 20 *B. tabaci* [42]. Therefore, raster values above 20 post-IDW were set to 20, and the resultant raster was normalised with a maximum value of 1.

The instantaneous state of the model is defined by the number of susceptible and infectious fields in each raster cell. The model is updated via a discrete event, continuous time stochastic process using an optimised Gillespie algorithm [35, 36]. Spatial coupling between infected and susceptible cells is governed by an isotropic discrete power law dispersal kernel, *K*, where the distance between the centroids of two raster cells *i* and *j* is *d*_*ij*_ and *α* is the exponent thus:

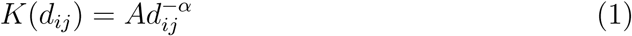

Due to an absence of data independently quantifying the two dispersal mechanisms, the model integrates both the net effects of dispersal by *B. tabaci* and the movement of planting material on the spread of infection into the single kernel.

An additional parameter, *p*, defines the kernel value at *d* = 0, where *p* is the proportion of dispersed inoculum that remains within the source cell. A kernel cut-off distance, *D*_*max*_ = 500km, sets the maximum distance from the source cell that the kernel covers. For the finite set of cell centroids in the range 0 *≤ d ≤ D*_*max*_, values are calculated based on the kernel function. A normalisation factor, *A*, is applied such that the sum of kernel values for *d >* 0 is equal to the value of 1 *−p*. Therefore, the sum of the kernel is 1.

The force of infection at location *i, ϕ*_*i*_, incorporates the kernel function, *K*, the transmission rate, *β*, the vector abundance parameter at location *j, w*_*j*_, and the current number of hosts in the infectious state, *I*_*j*_, where *j* represents all locations in the rasterised landscape, including *i* (Equation 2). The instantaneous rate of infection at a given raster cell, *i*, from all locations, *j*, is *ψ*_*i*_, that incorporates the vector abundance parameter, *w*_*i*_, and the number of susceptible hosts, *S*_*i*_, at location *i* (Equation 3). The effect of an infection event at location *i* is given by Equation 4. We assume a linear relationship between vector abundance and its effect on infection and susceptibility up to a field-level mean of 20 *B. tabaci* [42]. Exploratory analyses on alternative model structures are outlined in Supplementary Methods S1.1.3.

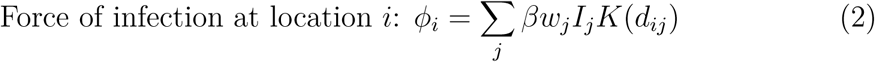

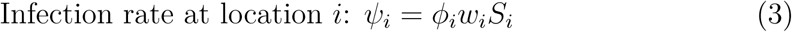

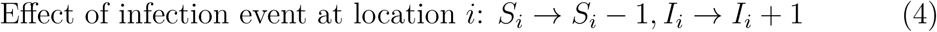

The model implements a surveillance scheme that replicates the real-world surveillance structure and intensity. For all years that surveillance was carried out in Uganda [14], we perform one instantaneous survey at the end of the simulation year. For example, we assume that all surveys that are carried out in 2005 are representative of the state on 31st December 2005. For each raster cell in the model landscape, we summed the number of fields that were surveyed in the Ugandan national survey within the bounds of the 1 km^2^ cell for a given survey year. We then randomly sampled the equivalent number of fields in each model cell and the numbers of sampled fields that were in each system state of susceptible and infectious were recorded allowing for the false negative survey detection rate of 0.15 (CBSD surveillance data in Methods).

The first reported observations of CBSD epidemic in Uganda occurred in November 2004 [13]. However, the study did not provide exact coordinate locations for the CBSD positive fields in November 2004. The dataset includes survey data from January 2005, reporting infected fields in the same region. Therefore, we assume the small number of CBSD infected fields reported during the January 2005 surveys is representative of the state of the epidemic on 1st January 2004, which we take as the simulation start time.

### 4.3 Parameter estimation

In the case of the Ugandan CBSD survey data, the model is unlikely to reproduce the spatiotemporal pattern of over 7600 records of CBSD presence/absence exactly. Therefore, given finite computational resources, the ABC methodology involved accepting parameter values from simulations that generated simulated survey data, *d*_*sim*_, that were considered sufficiently close to the real world data, *d*_*real*_. Summary statistics, *S*, were used to simplify the comparison between simulated and real data, along with the selection of a distance measure, *ρ*, and a maximum allowed distance (tolerance), *ϵ*, between *S*(*d*_*sim*_) and *S*(*d*_*real*_) defined:

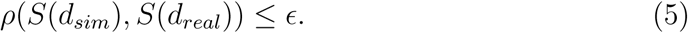

The target data for *S*_*cen*_ and *S*_*nat*_ were derived by applying the statistics to the real-world survey data, *S*_*cen*_(*d*_*real*_) and *S*_*nat*_(*d*_*real*_), resulting in the yearly proportion of survey sites within their geographical regions (the central area surrounding Kampala and the remaining area of Uganda not covered by the central area respectively) that were reported as positive for CBSD symptoms (Figure 2). The *S*_*inf*_ distance measure, *ρ*_*inf*_, is calculated by taking the maximum of the absolute annual differences between simulated and real survey proportions of infected survey sites (Equation 6). A tolerance, *ϵ*_*inf*_, then governs the maximum allowed value of *ρ*_*inf*_ (Equation 7).

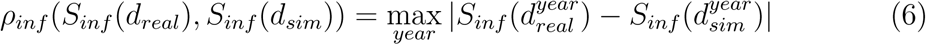

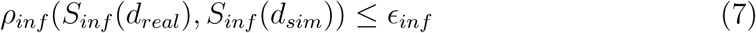

The statistic *S*_*grid*_ applied to the real Ugandan survey data, *d*_*real*_, has a perfect score of 1 (Equation 8). The distance measure, *ρ*_*grid*_ is calculated by subtracting the statistic as applied to simulated data, *d*_*sim*_, from 1 (Equation 9) and the tolerance governs the maximum allowed value of the distance measure (Equation 10). A deviation of *±*1 year for a given quadrat in the year in which CBSD was first detected in the Ugandan surveillance data is only allowed for quadrats with surveys both one year earlier and one year later than the target infection year, otherwise no deviation is allowed. Supplementary Figure 6 summarises the target data for each quadrat and highlights the quadrats with surveys both years either side of the target first year of infection.

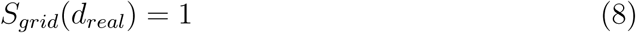

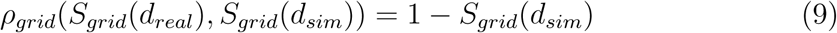

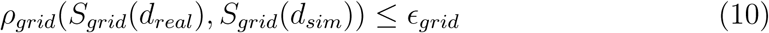

The summary statistic assessment methodology using synthetic data allowed an exploration of the convergence of the posterior distribution to the known parameter values as the tolerances are reduced, whilst retaining enough simulations to enable a smooth posterior distribution given the finite number of fitting simulations (Supplementary Methods S1.3). Based on these analyses, we use all three statistics in combination and define the fitting criteria as the following values for the three tolerances for each of the statistics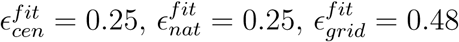

For the prior distribution of *p*, we sampled from a uniform distribution between 0 and 1. For *α* and *β*, we carried out multiple batches of simulations, updating the search space at each iteration to sufficiently explore parameter space in order to identify the regions of highest likelihood density (Supplementary Methods S1.2). The total number of fitting simulations was 233,600.

### 4.4 Model validation

We assess the ability of the parameterised model to predict the validation dataset, 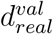, which spans 2011-2017. We run 10,000 validation simulations starting in 2004 using the same initial conditions as during parameter estimation and isolate the subset of simulations that meet the fitting criteria. We take the subset of simulations that pass the fitting criteria as representative of the system state at the start of 2011, then for the validation period calculate the summary statistics, *S*_*inf*_, and score model performance according to the percentage of simulations that satisfy 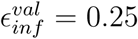. In addition, we present the full dynamics of *S*_*inf*_ (*d*_*real*_) and spatial comparisons of a simulated survey with the real-world survey.

### 4.5 Code availability

The code is available at https://github.com/camepidem/cbsd_model_development

### 4.6 Data availability

A script to automatically download the datasets analysed during this study from their published sources is shared as part of the code repository.

## 5 Acknowledgements

The authors gratefully acknowledge financial support from the Bill & Melinda Gates Foundation, the UK Foreign, Commonwealth and Development Office and the BB-SRC. We also acknowledge many helpful discussions and support from members of the Epidemiology & Modelling Group in Cambridge, especially Anna Szyniszewska and Renata Retkute, in addition to partners from the Cassava Diagnostics Project (CDP) and the Central and West African Virus Epidemiology (WAVE) project.

## 6 Author contributions

C.A.G., R.O.J.H.S. and D.G. formulated the problem and modelling approach. D.G. and R.O.J.H.S. developed, tested and implemented the modelling framework and performed parameter estimation and validation. T.A., P.A. and G.O. provided the surveillance data underpinning the analysis and gave expert insight into the clean seed programmes. D.G., C.A.G. and R.O.J.H.S planned and D.G. wrote the manuscript and created the figures in collaboration with C.A.G. and R.O.J.H.S.. C.A.G. supervised the project.

