## Supplementary information for "Developing a predictive model for an emerging epidemic on cassava in sub-Saharan Africa"

### S1 Supplementary Methods

#### S1.1 Model development

In order to develop a parameterised spatial model of the cassava brown streak disease epidemic, we began by considering a range of model structures. The models we considered included different dispersal kernels, the incorporation of data for vector abundance, and the incorporation of survey bias. For each model structure, we carried out preliminary model parameterisation via ABC rejection using a subset of the Ugandan CBSD surveillance data ($d_{real}^{fit}$) (see also Parameter estimation in Main Text Methods) and preliminary validation against the remaining surveillance data ($d_{real}^{val}$) (Model validation in Main Text Methods), which allowed us to quantify the predictive power of each model. In this context, the term preliminary simply means the number of fitting and validation simulations were lower than for the final model presented in the Main Text. These analyses led to the rejection of the use of an exponential dispersal kernel and the selection of the model structure showing the best correspondence to the validation surveillance data.

All models shared a set of foundational features. Firstly, as cassava is a vegetatively propagated crop, all models comprised an SI compartmental structure as infection persists from harvest to the subsequent planting [1]. Secondly, all models included the cassava host landscape (Incorporating data-driven model layers in Main Text

Methods). Parsimony guided our process of model development to find the simplest epidemiologically-informed model structure that can reproduce and predict the dynamics of the CBSD epidemic. Supplementary Figure 1 gives an overview of the parameterisation and validation methodology as applied during model development and for the final selected model.

#### S1.1.1 Estimating false negative survey detection rate

The surveillance protocol involved sampling 30 plants from only the dominant cassava variety in a field (CBSD surveillance data in Main Text Methods). For a subset of the Ugandan surveys, surveyors recorded whether CBSD symptoms were present anywhere in the field, not just from the 30 plant sample or the dominant cultivar. Therefore, we can use this subset of surveys to approximate the level of disease underestimation in the field. Supplementary Figure 2 shows the variability in this underestimation rate across the 5 years with an approximate mean of 0.15, which was taken as the simulated surveillance false negative rate.

#### S1.1.2 Kernel selection

During exploratory stages of model development, two candidate kernel functions were considered: an exponential (Equation S1) and a power law (Equation S2):

$$K(d) = Ae^{-\alpha d}, \tag{S1}$$

$$K(d) = Ad^{-\alpha}. \tag{S2}$$

An exponential kernel produces higher rates of local spread and lower rates of longer range spread. In contrast, a power law kernel has slightly higher rates of long range spread and lower rates of local spread. The CBSD spatiotemporal pattern observed in the surveillance data from Uganda indicated significant long range spread combined with lower rates of local bulk up [2]. Preliminary simulations confirmed that an exponential kernel showed poor correspondence with the data (results not presented) compared with a power law kernel.

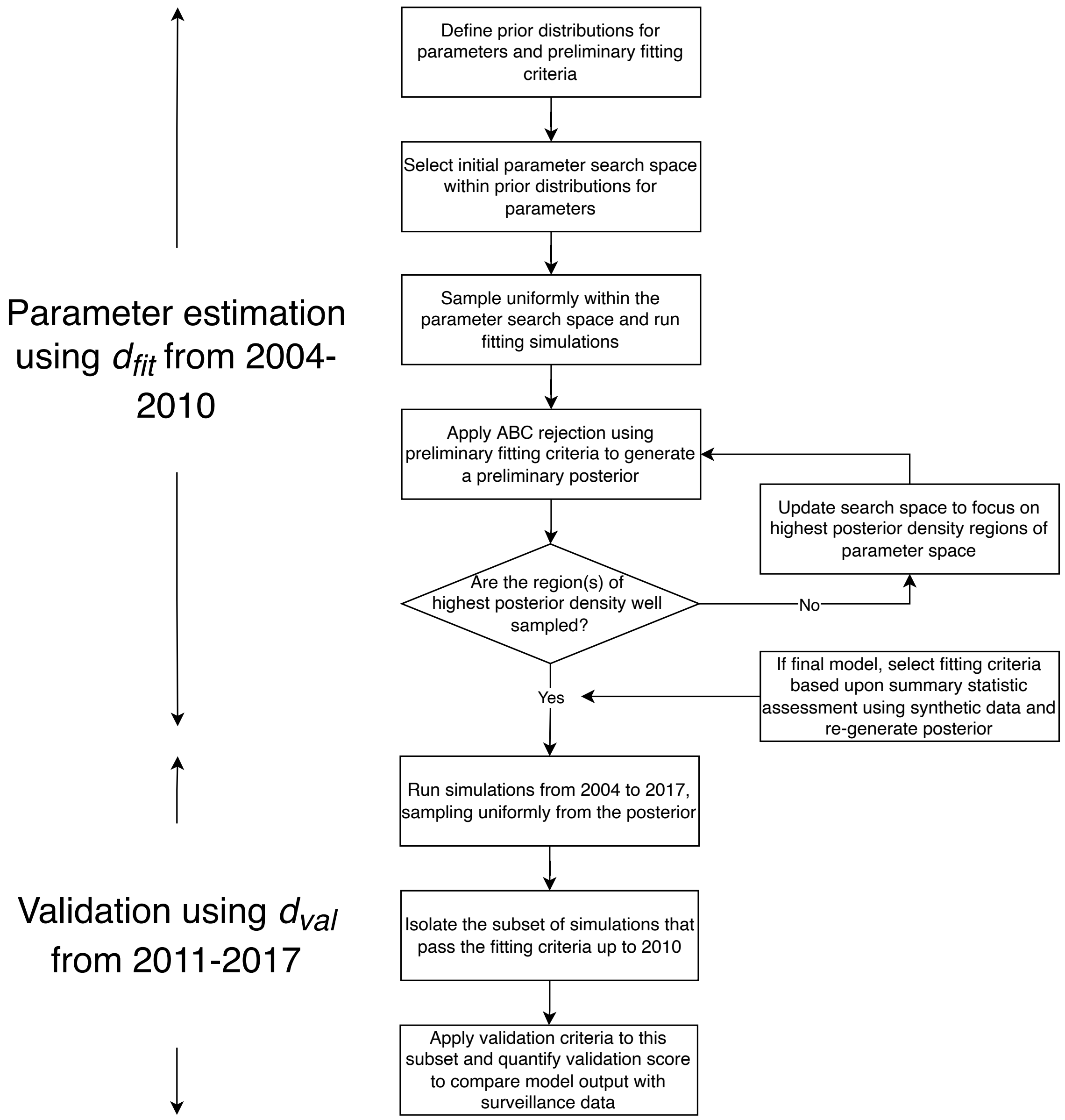


Supplementary Figure 1: Overview of the parameterisation and validation methodology as applied during model development. Candidate models are proposed and preliminary fitting and validation is carried out to assess the ability of the model to reproduce the dynamics of $d^{fit}_{real}$ and predict the dynamics of $d^{val}_{real}$. For the candidate model that performs best in comparison with the other candidates during preliminary validation, additional batches of simulations are run covering the broad region of highest posterior density to improve the sampling density. The summary statistic assessment is then carried out to identify appropriate summary statistic tolerances for the fitting criteria, given the density of sampling from the prior.

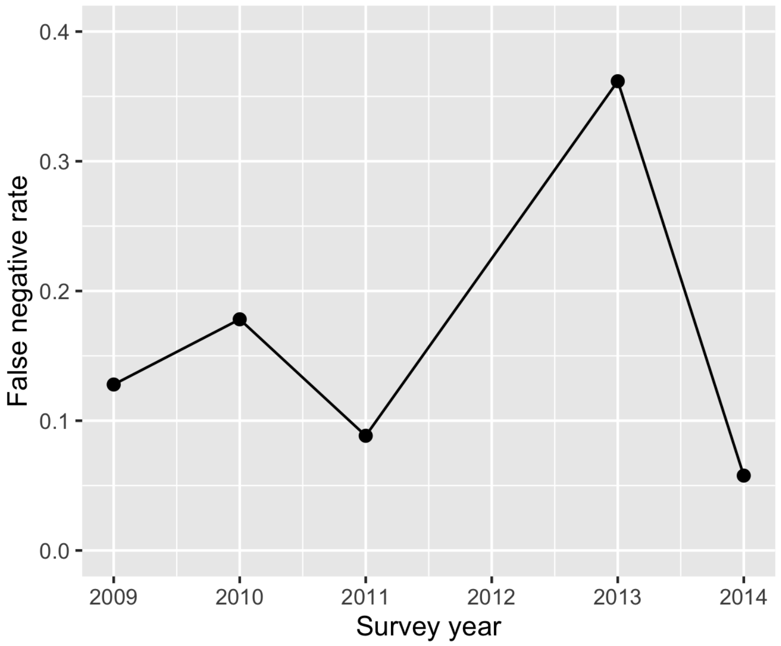


Supplementary Figure 2: Variability across survey years in the rate at which a field is reported as not having any CBSD foliar symptoms in the 30 plant sample of the dominant cultivar, whereas CBSD foliar symptoms were in fact present on the non-dominant cultivar(s) or beyond the 30 plant sample. The rates are estimated from additional information provided about CBSD observations for a subset of surveys in the Ugandan CBSD dataset [2].

| Model index | False negative allowance | Vector abundance |
|---|---|---|
| 0 | ✗ | ✗ |
| 1 | ✓ | ✗ |
| 2 | ✓ | ✓ |

Supplementary Table 1: Model features added to the base model framework for the three candidate models.

#### S1.1.3 Candidate models

We iteratively developed and parameterised three candidate models based on the selection of the power law kernel, which we refer to as model_0, model_1 and model_2. The simplest model comprised the host landscape and the SI structure with a power law kernel. The two subsequent models built upon this foundation. We assessed the performance of the initial model (model_0), then iteratively added additional features. Supplementary Table 1 summarises the features included in each model iteration.

The two additional features included in model iterations model_1 and model_2 were the allowance for a false negative survey detection rate and the spatial vector abundance layer. The false negative survey detection rate refers to the modification of the simulated surveillance scheme to allow for false negatives in cases, such as surveyors not recording disease when present on varieties other than on the dominant variety in a field (cf. CBSD surveillance data in Main Text Methods and Supplementary Methods S1.1.1). The spatial vector abundance layer introduced in model_2 represents the relative abundance of the CBSI vector, *B. tabaci*, at a 5 km resolution. The vector layer allowed the rate at which infection spreads to, from, and within a given location to be modulated by the local abundance of the vector (Figure 1 and see also Model structure in Main Text Methods).

For each model, we carried out preliminary ABC parameter estimation and model validation. In contrast to the method described for the final model in the Main Text, during model development the number of fitting simulations was limited to between 20,000 to 30,000 and the number of validation simulations to 1,000. The three parameters to be estimated were the proportion of dispersal that remains within the source cell, $p$, the kernel scale parameter, $\alpha$, and the transmission rate, $\beta$. Supplementary Figure 3 shows the posterior distributions obtained from preliminary parameter estimation for each of the three models. The tolerances applied to the three summary statistics were $\epsilon^{fit}_{inf} = 0.3$, $\epsilon^{fit}_{grid} = 0.58$, which we define as the

| Model index | No. simulations | $P(d_{real}^{fit})$ | $P(d_{real}^{fit} \cap d_{real}^{val})/P(d_{real}^{fit})$ |
|---|---|---|---|
| 0 | 1000 | 0.24 | 0.004 |
| 1 | 1000 | 0.27 | 0.87 |
| 2 | 1000 | 0.36 | 0.98 |

Supplementary Table 2: Validation performance of each model. $P(d_{real}^{fit})$ refers to the proportion of the simulations that pass within the tolerances applied to the summary statistics for the fitting period. $P(d_{real}^{fit} \cap d_{real}^{val})/P(d_{real}^{fit})$ refers to the proportion of simulations that, on a per simulation basis, pass within both the fitting and validation tolerances, divided by the proportion that pass within just the fitting tolerances.

preliminary fitting criteria.

Supplementary Figure 4 shows the behaviour of each model from an ensemble of $1,000$ validation simulations through the lens of the two infectious proportion-based metrics. All three models appear capable of reproducing the infectious proportion dynamics in the central area surrounding Kampala. However, there is a clear improvement in performance for the Ugandan national infectious proportion metric from model_0 to model_2. In order to quantify the performance of the preliminary validation simulations, the same methodology as described in the Main Text is used, but with different tolerances. Specifically, for each 1,000 validation simulations for each model, we calculate the proportion of simulations that meet the preliminary fitting criteria. For this subset of simulations, we calculate the proportion that sufficiently correspond to the validation dataset, $d_{real}^{val}$, which we define as $\epsilon_{inf}^{val} = 0.3$. Supplementary Table 2 summarises these results. The strong preliminary results of model_2 justified our selection of this model structure for a final round of more intensive ABC parameter estimation.

##### S1.1.4 Estimating the clean seed replacement parameter

The parameterised version of model_2 is used to estimate the value of the clean seed replacement proportion parameter, $r_{clean}$ (cf. Parameter estimation in Main Text Methods). A parameter sweep covered values of $r_{clean}$ from 0 to 0.4 in increments of 0.05, with an ensemble of 1000 simulations per parameter value. We then applied the validation criteria (cf. Model validation in Main Text Methods) but with modified tighter tolerances of $\epsilon_{val}^{cen}$ of 0.15 across the clean seed programme period of 2013-2015 to differentiate the comparative performance of each value of $r_{clean}$. The tolerance was retained as $\epsilon_{val}^{cen} = 0.25$ for validation years outside of the clean seed programme

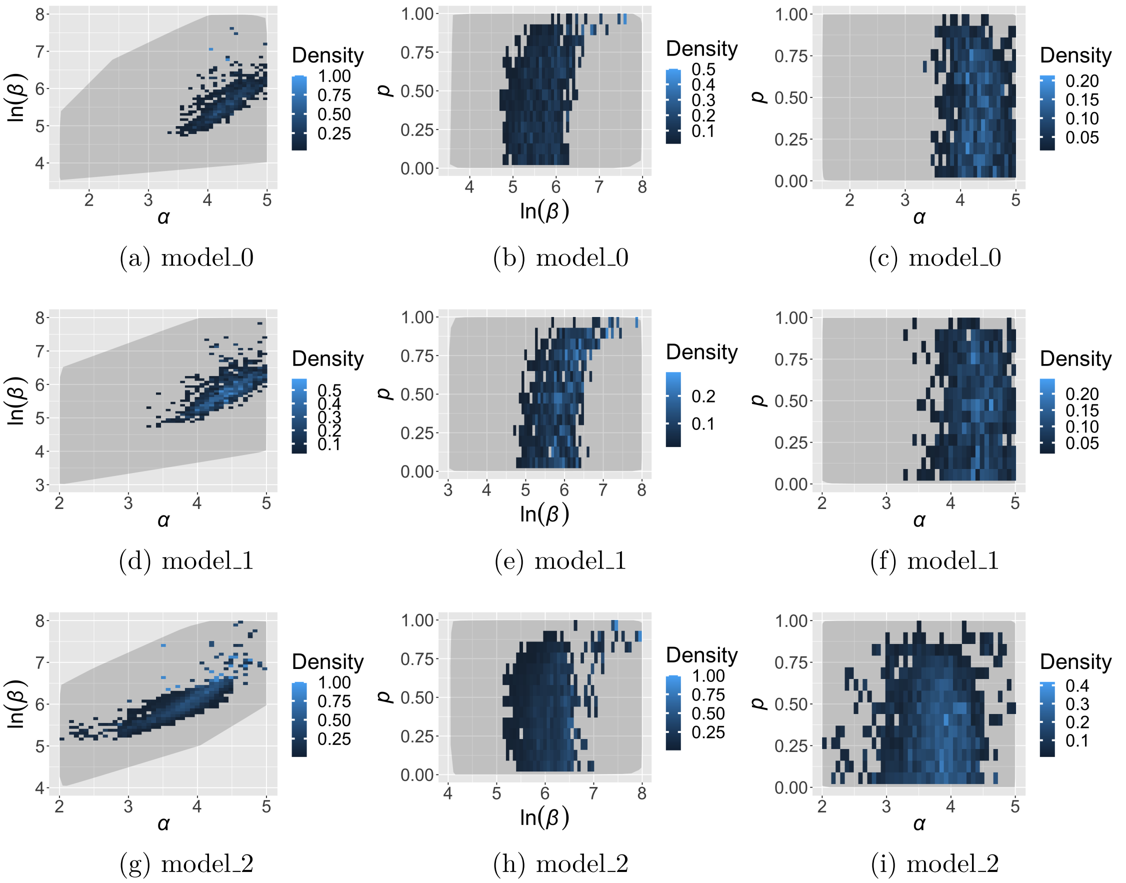


(a) model_0 (b) model_0 (c) model_0

(d) model_1 (e) model_1 (f) model_1

(g) model_2 (h) model_2 (i) model_2

Supplementary Figure 3: Posterior distributions selected for preliminary model validation for each of the three models. (a) to (c): model_0; (d) to (f): model_1; (g) to (i): model_2.

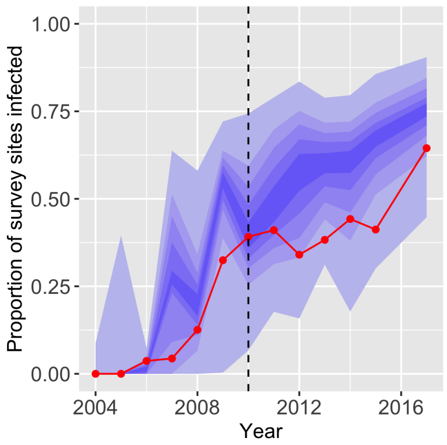


(a) model_0 - $S_{nat}$

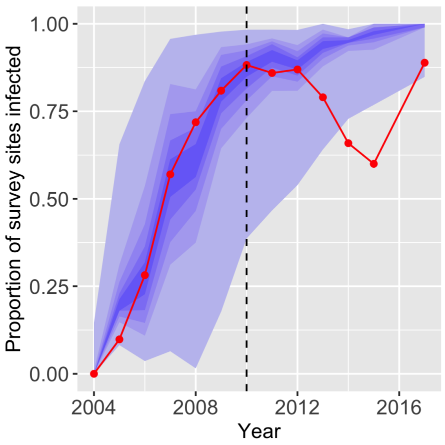


(b) model_0 - $S_{cen}$

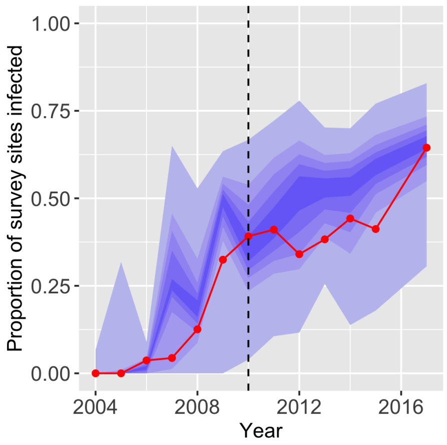


(c) model_1 - $S_{nat}$

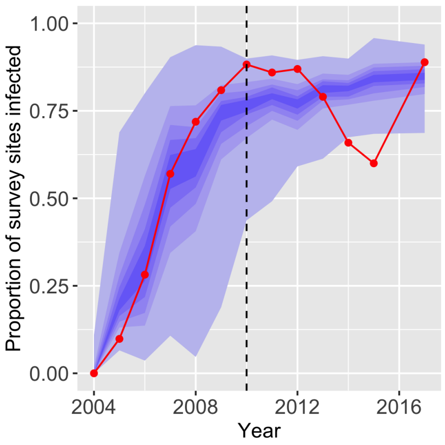


(d) model_1 - $S_{cen}$

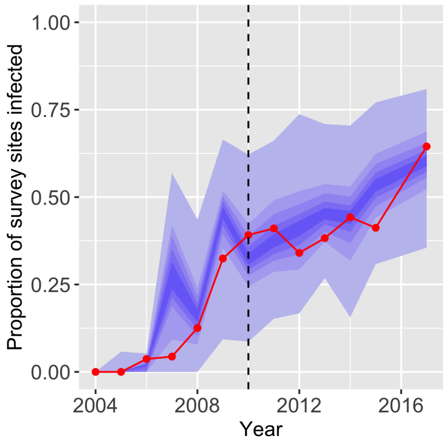


(e) model_2 - $S_{nat}$

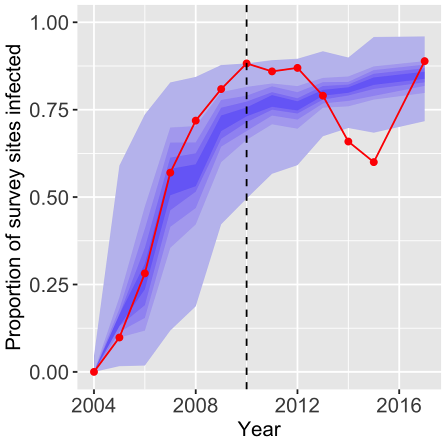


(f) model_2 - $S_{cen}$

Supplementary Figure 4: Comparison of model dynamics across the three model structures through the lens of the two summary statistics, $S_{nat}$ and $S_{cen}$, based on 1000 validation simulations from each models' respective posterior distributions and without applying any tolerances. All models appear to perform similarly based on $S_{cen}$ but $S_{nat}$ improves across model iterations. The red line indicates the target value for the statistic derived from surveillance data. The central blue band is the median $\pm 10\%$ and each gradation beyond is a further $\pm 10\%$ from the median.

| $r_{clean}$ | No. simulations | $P(d^{fit}_{real})$ | $P(d^{fit}_{real} \cap d^{val}_{real})/P(d^{fit}_{real})$ |
|---|---|---|---|
| 0 | 1000 | 0.008 | 0 |
| 0.05 | 1000 | 0.003 | 0.33 |
| 0.1 | 1000 | 0.007 | 0.57 |
| 0.15 | 1000 | 0.013 | 0.92 |
| 0.2 | 1000 | 0.004 | 0.75 |
| 0.25 | 1000 | 0.008 | 0.125 |
| 0.3 | 1000 | 0.010 | 0.1 |
| 0.35 | 1000 | 0.005 | 0 |
| 0.4 | 1000 | 0.010 | 0 |

Supplementary Table 3: Results of the parameter sweep of $r_{clean}$, the proportion of CBSI infected fields replaced with clean planting material for each of the three yearly rounds of intervention from 2013 to 2015 in four districts of Uganda. For the ensemble of simulations with each value of $r_{clean}$, $P(d^{fit}_{real})$ refers to the proportion of simulations that pass within the tolerances applied to the summary statistics for the fitting period. $P(d^{fit}_{real} \cap d^{val}_{real})/P(d^{fit}_{real})$ refers to the proportion of simulations that, on a per simulation basis, pass within both the fitting and validation tolerances, divided by the proportion that pass within just the fitting tolerances. Unlike model validation in the main text, where $\epsilon^{cen}_{val}$ is fixed at 0.25, in this case, a tighter tolerance value of $\epsilon^{cen}_{val}$ of 0.15 was applied during the period of 2013 to 2015 to more effectively differentiate the performance of the model with each value of $r_{clean}$.

period. Supplementary Table 3 summarises the proportion of simulations for each value of $r_{clean}$ that pass the fitting and validation criteria with the tighter tolerances and indicates that a value of 0.15 for $r_{clean}$ best captures the dynamics of the clean seed programme (Supplementary Figure 5).

### S1.2 Parameter estimation

#### S1.2.1 Efficient sampling from the prior

For the process of ABC rejection applied to computationally intensive models, a major challenge is to run enough simulations to explore the prior sufficiently with finite computational resources [3]. We used a high performance computing (HPC) cluster, which allowed us to parallelise the fitting and validation simulations. Nonetheless, the total compute time for the 233,600 fitting simulations was 87,157 hours, with a mean simulation time of 23 minutes. Therefore, in order to make efficient use of computational resources, for parameters $\alpha$ and $\beta$, we sampled from a joint uniform prior and carried out multiple batches of simulations, updating the search space at

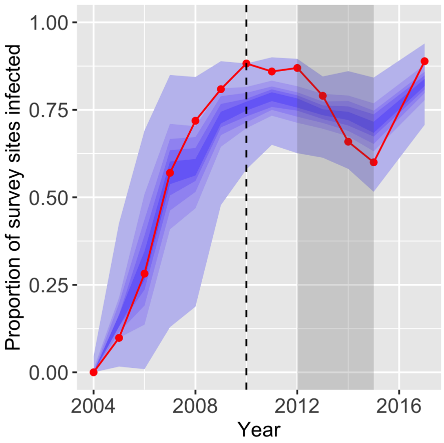


(a) $r_{clean} = 0.1$

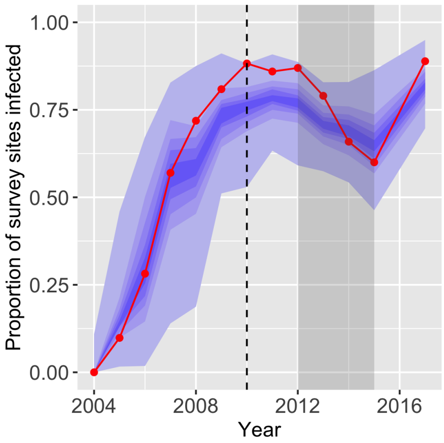


(b) $r_{clean} = 0.15$

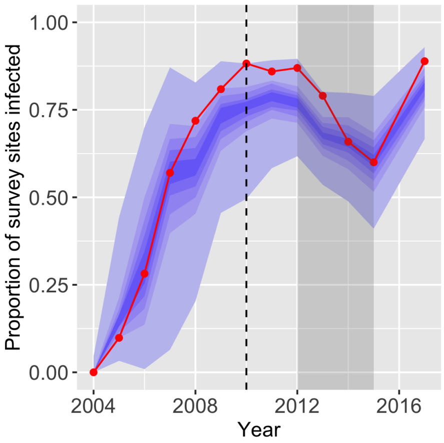


(c) $r_{clean} = 0.2$

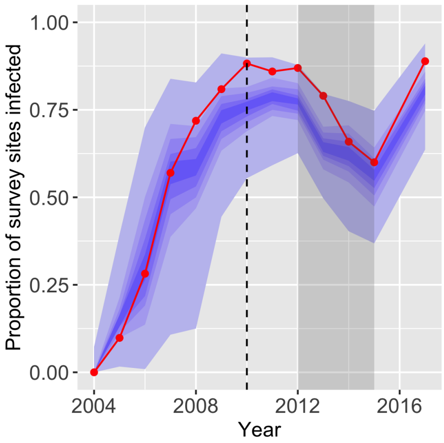


(d) $r_{clean} = 0.25$

Supplementary Figure 5: A comparison of the impact of different values of $r_{clean}$ on $S_{cen}$. The grey band indicates the period in which we implemented a process equivalent to the clean seed programmes via three discrete rounds of $I \rightarrow S$ replacement at the start of the 2013, 2014 and 2015 growing seasons. The red line indicates the statistic applied to the real-world surveillance data, $S_{cen}(d_{real})$. The black vertical dotted line indicates the boundary between $d_{real}^{fit}$ and $d_{real}^{val}$. A quantitative assessment indicated a value of 0.15 best captures the impact of the clean seed programme on $S_{cen}$ (Supplementary Table 3).

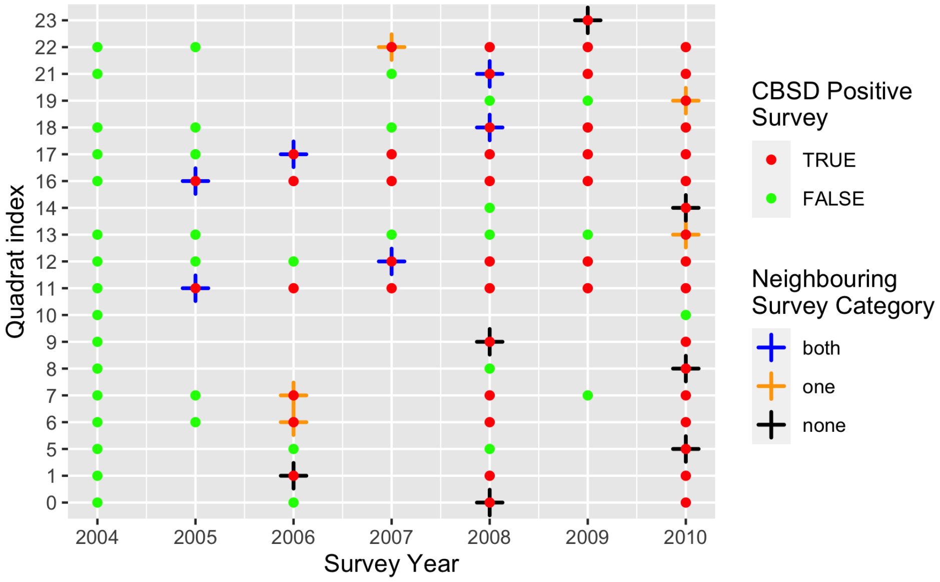


Supplementary Figure 6: Summary of the timecourse of surveys for each quadrat and whether or not CBSD was reported from any surveyed fields in each year up to 2010. In addition, for quadrats in which CBSD is detected, crosses indicate the first year of CBSD reporting and are shaded according to whether surveys were carried out ±1 year either side of the year of first detection.

each iteration (Supplementary Figure 7). For the proportion of dispersed inoculum that remains in the source cell, $p$, the search space remained the full uniform prior between 0 and 1. For all parameters, we sampled continuously within parameter space without discretisation.

Carrying out sequential batches in this way is a more efficient use of resources, allowing a wide initial exploration of parameter space whilst spending the majority of computational resource on regions of highest likelihood density (i.e. more relevant regions). Initially sampling from the full search space defined by the prior (see batch_0 of Supplementary Figure 7) ensures that we identify the optimal region(s) before restricting the search space to regions of higher likelihood density. Moreover, this allows us to apply the method that uses the summary statistics to recover known parameters using synthetic data, to ensure that the fitting techniques can successfully discern between parameters from different regions of parameter space (Supplementary Methods S1.3).

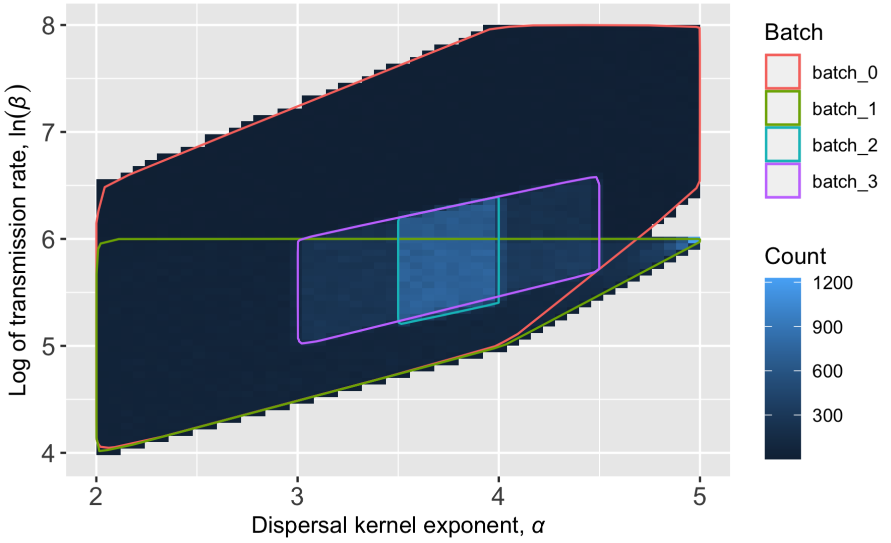


Supplementary Figure 7: The density of sampling of the kernel exponent, $\alpha$, and the log of the transmission rate, $\ln(\beta)$, during parameter estimation. A uniform prior was used for the third estimated parameter, $p$, being the proportion of dispersed inoculum that remains in the source cell. Simulations were carried out in batches, iteratively exploring and updating the parameter search space for each subsequent batch, with the aim of converging towards and densely sampling the region with highest posterior probability density. The total number of simulations in batches 0 to 3 respectively are: 22,603, 71,183, 59,878, 79,936.

##### S1.2.2 Calculating the posterior distribution

For each of the estimated parameters, we discretise parameter space into cuboids of the following dimensions: $\alpha$: 0.02, $\ln(\beta)$: 0.1, $p$: 0.02. The approximate likelihood is calculated as the number of accepted samples within each cuboid divided by the total number of samples within the cuboid. Note that the number of samples per cuboid affects the width of the confidence interval obtained for likelihood values, not the likelihood itself. We choose to average samples over cuboids, rather that taking individual samples as it allows pooling of many samples to get a smoother posterior distribution, minimising noise in the final output.

As the prior is uniform, the approximate likelihood is taken as the posterior. When simulating using the posterior, we sample from the distribution of cuboids, first selecting a cuboid according to it's posterior value and then selecting parameter values by sampling uniformly within the selected cuboid.

### S1.3 Summary statistic assessment using synthetic data

The goal of estimating parameters using the Ugandan CBSD surveillance data is to calibrate the model to the dynamics of the spatio-temporal spread observed in the data. Therefore, the process of summary statistic selection is to identify statistics with the ability to capture the landscape-scale spatiotemporal spread behaviour of the CBSD epidemic. However, the optimal method for selecting summary statistics and tolerances remains an open question in the likelihood-free inference literature [4–6]. For high dimensional data, it may not be computationally feasible to use summary statistics that meet the statistical definition of sufficiency and it is common practice to select summary statistics based on their ability to capture an important property of the system dynamics, which are described as informative summary statistics. [7].

In the Main Text Methods we describe three informative statistics used for ABC parameter estimation. These statistics have been designed to discriminate between epidemics with different landscape-scale dynamic behaviour. Here, we outline the methods used to assess the suitability of a given statistic.

#### S1.3.1 Methods

The process of summary statistic assessment begins by selecting model parameters from the prior distribution and running model_2 using these known parameters to generate synthetic epidemic surveillance data, $d_{syn}$. The location and timing of simulated surveillance for the synthetic data is retained from the real-world Ugandan surveillance data.

We then perform the ABC parameter estimation procedure (cf. Parameter estimation in Main Text Methods) to derive a posterior distribution for the parameters using a given statistic, $S$, or combination of statistics. Specifically, for each of the generated $d_{syn}$ data sets, ABC parameter estimation is carried out multiple times, first using each of the summary statistics, $S_{cen}, S_{nat}, S_{grid}$ independently, then using all three statistics in combination. For a given statistic or statistics in combination, as the tolerance(s) is reduced, $\epsilon \to 0$, we assess the behaviour of the statistic to discern between different regions of parameter space and converge towards the known input parameters.

| Param set | $\alpha$ | $\beta$ | $\ln(\beta)$ | $p$ | $n_{rep}$ | $n_{fit}$ |
|---|---|---|---|---|---|---|
| 0 | 3.5 | 403.43 | 6 | 0.5 | 10 | 233,600 |
| 1 | 3 | 121.51 | 4.8 | 0.2 | 10 | 233,600 |
| 2 | 4.5 | 1096.63 | 7 | 0.8 | 10 | 233,600 |

Supplementary Table 4: Input parameter sets selected for the generation of synthetic surveillance data, $d_{syn}$. For each parameter set, model_2 is run $n_{rep}$ times to generate $d_{syn}$ with stochastic variability. Model parameters are $\alpha$, the dispersal kernel exponent, $\beta$, the transmission rate and $p$, the proportion of dispersed inoculum that remains in the source cell. The same ensemble of fitting simulations, $n_{fit}$, used for model_2 parameter estimation with the real-world Ugandan survey data (see Main Text) is used for parameter estimation using $d_{syn}$.

To test whether a statistic is robust to the stochasticity of the model, the statistic evaluation procedure is repeated to generate 10 distinct sets of synthetic surveillance data for a given set of input parameters. By fitting to synthetic data, the selection procedure emulates the process of estimating parameters on the real-world data, but when the true parameters are known.

The entire process was repeated for three different parameter sets to generate synthetic data with different spread behaviour. The input parameter value sets are given in Supplementary Table 4. The proposed summary statistics are used independently and in combination to attempt to recover the known parameter values, hence assessing the robustness of the summary statistics to a diversity of epidemics.

#### S1.3.2 Results and discussion

For all of the presented results, we have selected representative figures from the 10 replicate simulations per parameter set. Supplementary Figure 8 shows the outputs of the simulated surveillance scheme from a representative synthetic simulation for each input parameter set. Importantly, each parameter set results in epidemics with visually distinct spread patterns.

Supplementary Figure 9 to Supplementary Figure 11 represent pairwise plots of the posterior distributions based on $\alpha$ and $\ln(\beta)$, derived by carrying out ABC parameter estimation using each of the summary statistics in isolation with a given $d_{syn}$ dataset as generated by the corresponding known parameters. The figures contrast posterior distributions derived using increasingly smaller tolerances. As the tolerance are reduced, the posterior distributions consistently converge towards the simulation parameters.

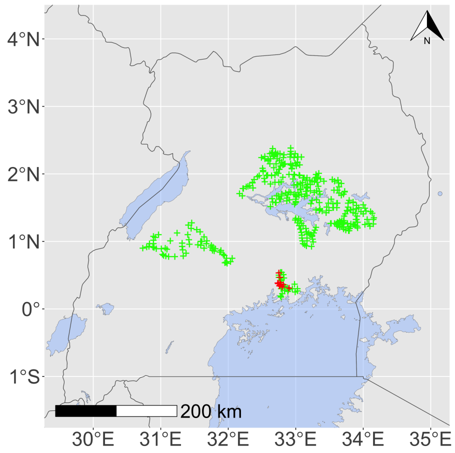


(a) Parameter set 0: 2005

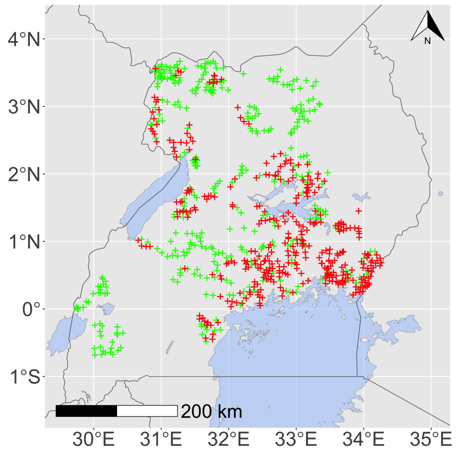


(b) Parameter set 0: 2010

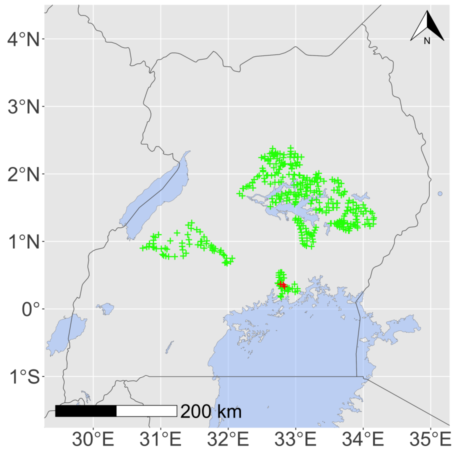


(c) Parameter set 1: 2005

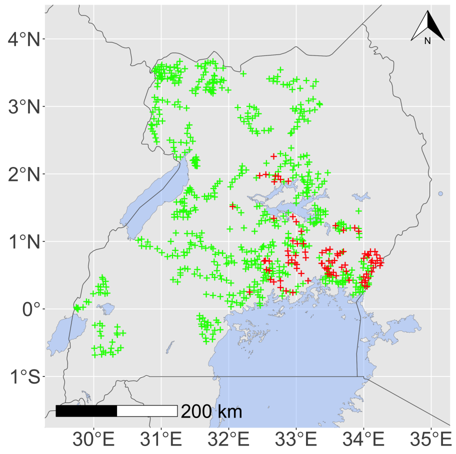


(d) Parameter set 1: 2010

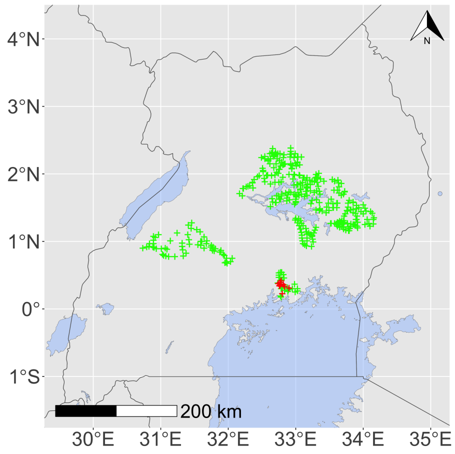


(e) Parameter set 2: 2005

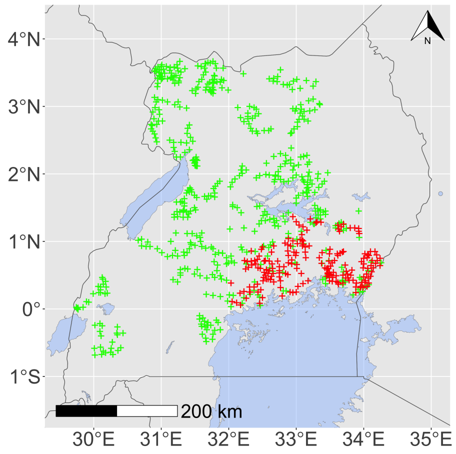


(f) Parameter set 2: 2010

Supplementary Figure 8: Synthetic surveillance data from a single realisation of each parameter set from Supplementary Table 4, highlighting the fundamental differences in the spatial structure of the different epidemics as the input parameters are varied. Red crosses indicated CBSD detection at the field level, and green crosses for a report of CBSD absence.

Supplementary Figure 12 illustrates the equivalent process using all three summary statistics in combination for ABC parameter estimation, which appears to perform especially well. Similarly, Supplementary Figure 13 and Supplementary Figure 14 show pairwise plots of the remaining parameter combinations when summary statistics are used in combination and also show a general trend of convergence to the known parameter values. The strong performance of the three statistics in combination motivated their selection for ABC parameter estimation with the Ugandan surveillance data.

It is important to note that a number of the pairwise posterior plots appear to be sparsely populated as the tolerances are reduced to very small values. This is due to non-uniform sampling density, wherein the majority of the fitting simulations are concentrated in the region of parameter space containing the highest posterior density for the real-world Ugandan survey data (Main Text Figure 3). Nonetheless, for each of the input parameter sets, the sampling intensity appears sufficient to indicate convergence towards the known parameter values.

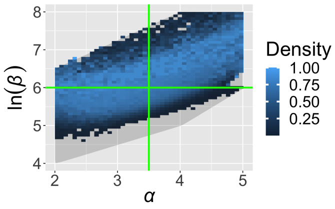


(a) param_0: 0.5 tol

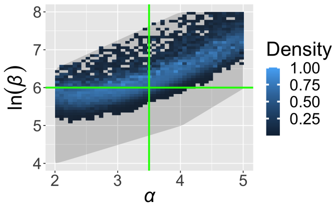


(b) param_0: 0.25 tol

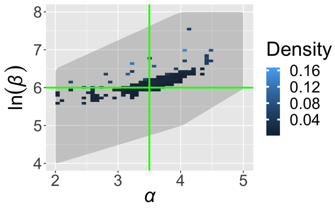


(c) param_0: 0.05 tol

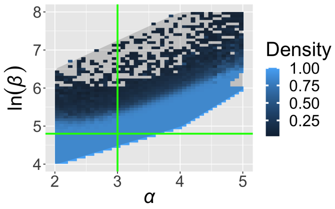


(d) param_1: 0.5 tol

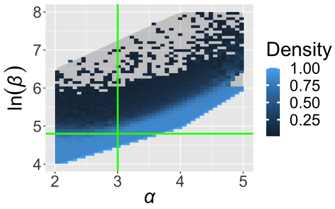


(e) param_1: 0.25 tol

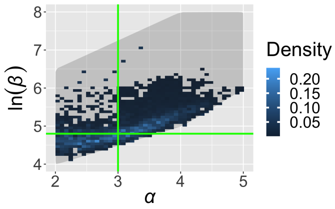


(f) param_1: 0.05 tol

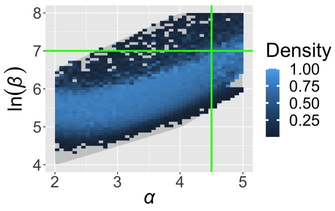


(g) param_2: 0.5 tol

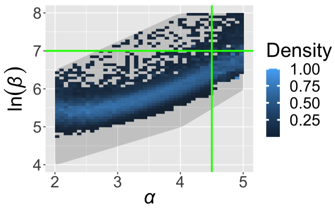


(h) param_2: 0.25 tol

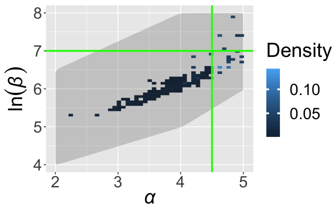


(i) param_2: 0.05 tol

Supplementary Figure 9: Recovering simulation parameters for the three different input parameter sets using $S_{cen}$, summarised as pairwise plots of $\alpha$ and $\ln(\beta)$. For each parameter set, $\epsilon_{cen}$ is reduced to inspect the convergence to the known parameter values. Target parameters are indicated by green lines. (a-c) is parameter set 0, (d-f) is parameter set 1, (g-i) is parameter set 2.

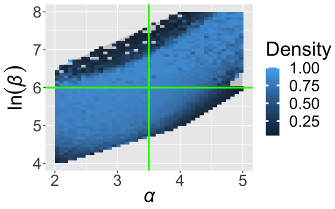


(a) param_0: 0.5 tol

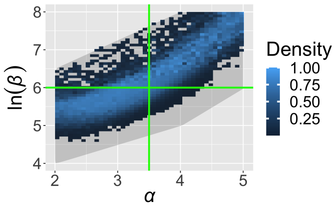


(b) param_0: 0.25 tol

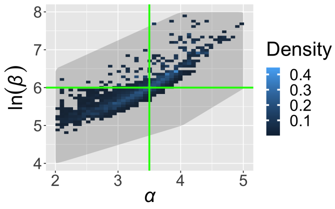


(c) param_0: 0.05 tol

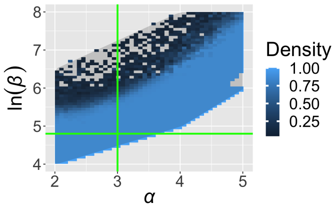


(d) param_1: 0.5 tol

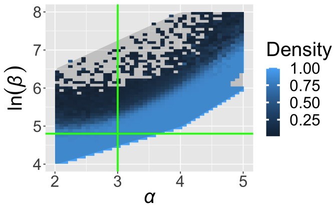


(e) param_1: 0.25 tol

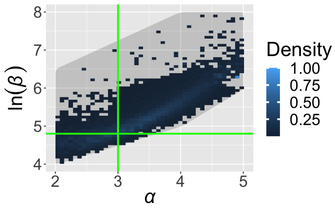


(f) param_1: 0.05 tol

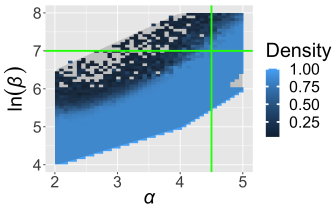


(g) param_2: 0.5 tol

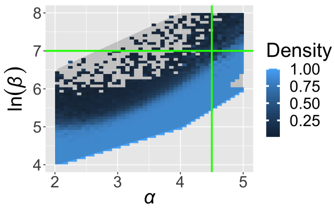


(h) param_2: 0.25 tol

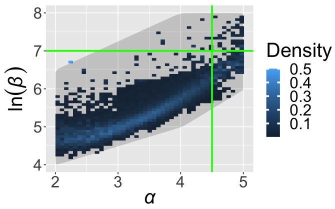


(i) param_2: 0.05 tol

Supplementary Figure 10: Recovering simulation parameters for three different input parameter sets using $S_{nat}$, summarised as pairwise plots of $\alpha$ and $\ln(\beta)$. For each parameter set, $\epsilon_{nat}$ is reduced to inspect the convergence to the known parameter values. Target parameters are indicated by green lines. (a-c) is parameter set 0, (d-f) is parameter set 1, (g-i) is parameter set 2.

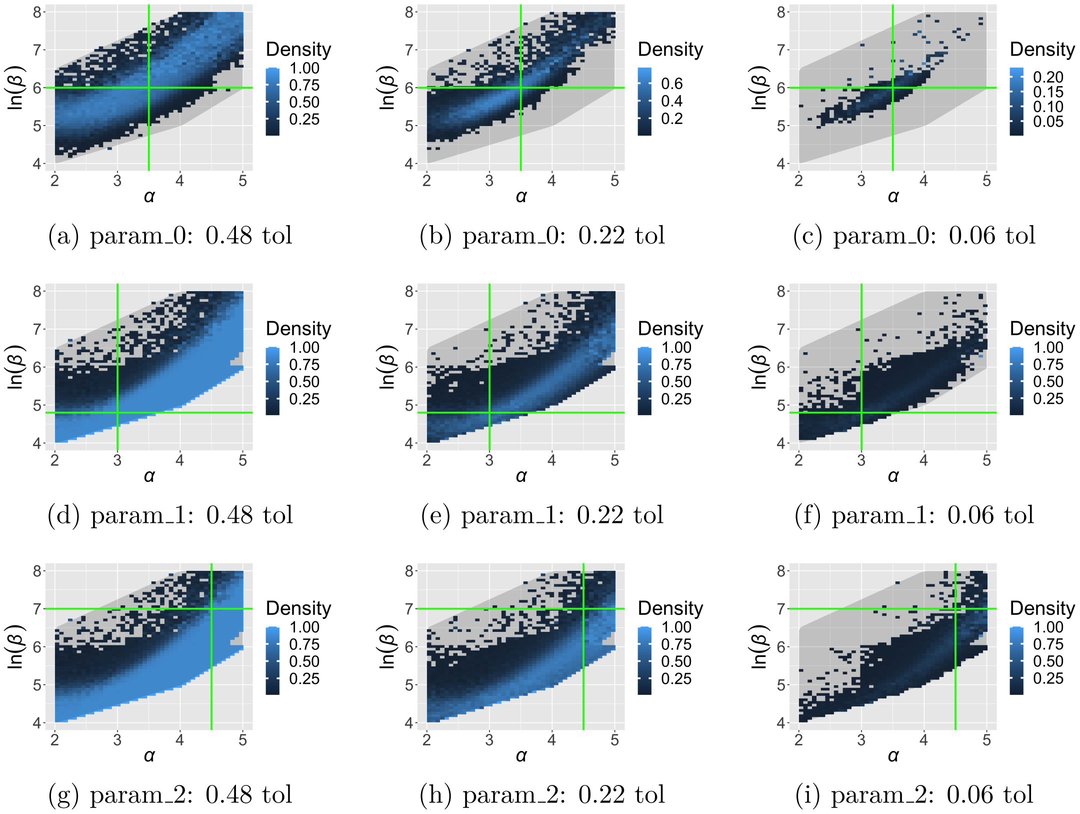


(a) param_0: 0.48 tol (b) param_0: 0.22 tol (c) param_0: 0.06 tol

(d) param_1: 0.48 tol (e) param_1: 0.22 tol (f) param_1: 0.06 tol

(g) param_2: 0.48 tol (h) param_2: 0.22 tol (i) param_2: 0.06 tol

Supplementary Figure 11: Recovering simulation parameters for three different input parameter sets using $S_{grid}$, summarised as pairwise plots of $\alpha$ and $\ln(\beta)$. For each parameter set, $\epsilon_{grid}$ is reduced to inspect the convergence to the known parameter values. Target parameters are indicated by green lines. (a-c) is parameter set 0, (d-f) is parameter set 1, (g-i) is parameter set 2.

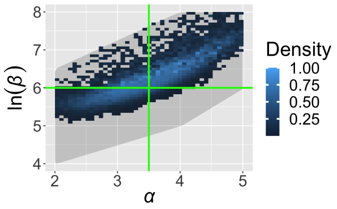


(a) param_0: 0.48 $\epsilon_{grid}$, 0.25 $\epsilon_{inf}$

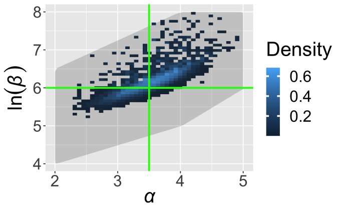


(b) param_0: 0.37 $\epsilon_{grid}$, 0.15 $\epsilon_{inf}$

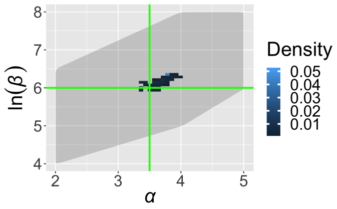


(c) param_0: 0.27 $\epsilon_{grid}$, 0.05 $\epsilon_{inf}$

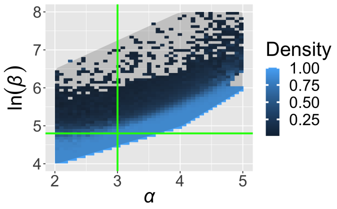


(d) param_1: 0.48 $\epsilon_{grid}$, 0.25 $\epsilon_{inf}$

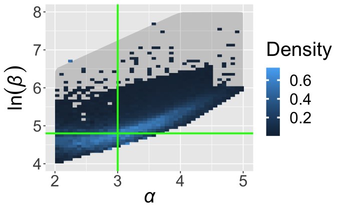


(e) param_1: 0.37 $\epsilon_{grid}$, 0.15 $\epsilon_{inf}$

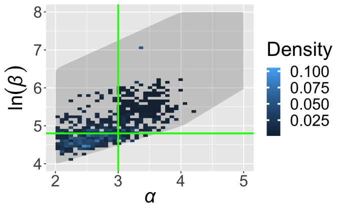


(f) param_1: 0.27 $\epsilon_{grid}$, 0.05 $\epsilon_{inf}$

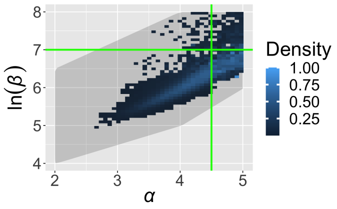


(g) param_2: 0.48 $\epsilon_{grid}$, 0.25 $\epsilon_{inf}$

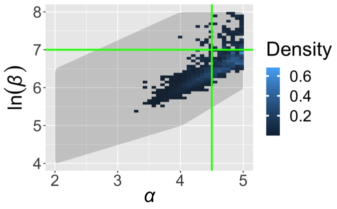


(h) param_2: 0.37 $\epsilon_{grid}$, 0.15 $\epsilon_{inf}$

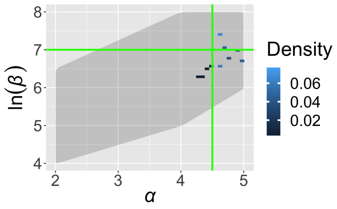


(i) param_2: 0.27 $\epsilon_{grid}$, 0.05 $\epsilon_{inf}$

Supplementary Figure 12: Recovering simulation parameters for three different input parameter sets using the three summary statistics in combination, $S_{cen}, S_{nat}, S_{grid}$, summarised as pairwise plots of $\alpha$ and $\ln(\beta)$. The tolerances for both of the infectious proportion-based summary statistics, $\epsilon_{cen}, \epsilon_{nat}$, are represented by $\epsilon_{inf}$. (a-c) is parameter set 0, (d-f) is parameter set 1, (g-i) is parameter set 2.

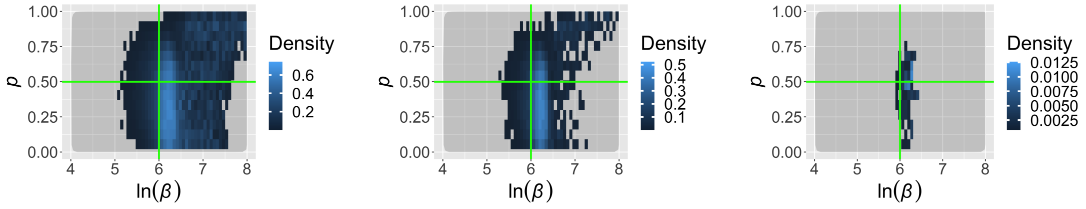


(a) param_0: 0.48 $\epsilon_{grid}$, 0.25 $\epsilon_{inf}$ (b) param_0: 0.37 $\epsilon_{grid}$, 0.15 $\epsilon_{inf}$ (c) param_0: 0.27 $\epsilon_{grid}$, 0.05 $\epsilon_{inf}$

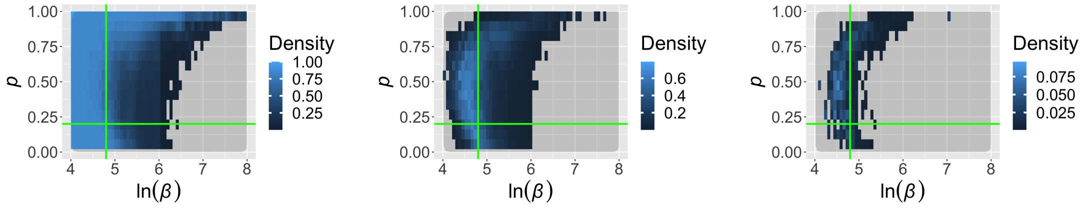


(d) param_1: 0.48 $\epsilon_{grid}$, 0.25 $\epsilon_{inf}$ (e) param_1: 0.37 $\epsilon_{grid}$, 0.15 $\epsilon_{inf}$ (f) param_1: 0.27 $\epsilon_{grid}$, 0.05 $\epsilon_{inf}$

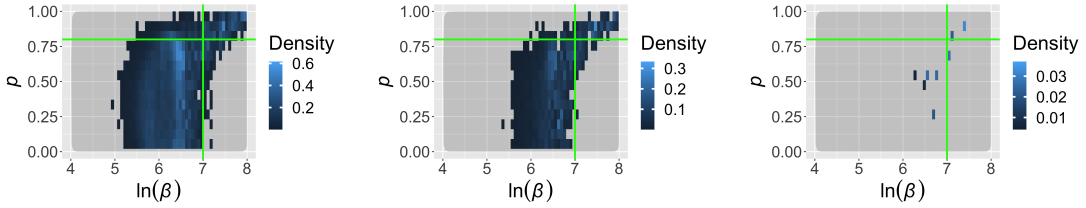


(g) param_2: 0.48 $\epsilon_{grid}$, 0.25 $\epsilon_{inf}$ (h) param_2: 0.37 $\epsilon_{grid}$, 0.15 $\epsilon_{inf}$ (i) param_2: 0.27 $\epsilon_{grid}$, 0.05 $\epsilon_{inf}$

Supplementary Figure 13: Recovering simulation parameters for three different input parameter sets using the three summary statistics in combination, $S_{cen}, S_{nat}, S_{grid}$, summarised as pairwise plots of $p$ and $\ln(\beta)$. The tolerances for both of the infectious proportion-based summary statistics, $\epsilon_{cen}, \epsilon_{nat}$, are represented by $\epsilon_{inf}$. (a-c) is parameter set 0, (d-f) is parameter set 1, (g-i) is parameter set 2.

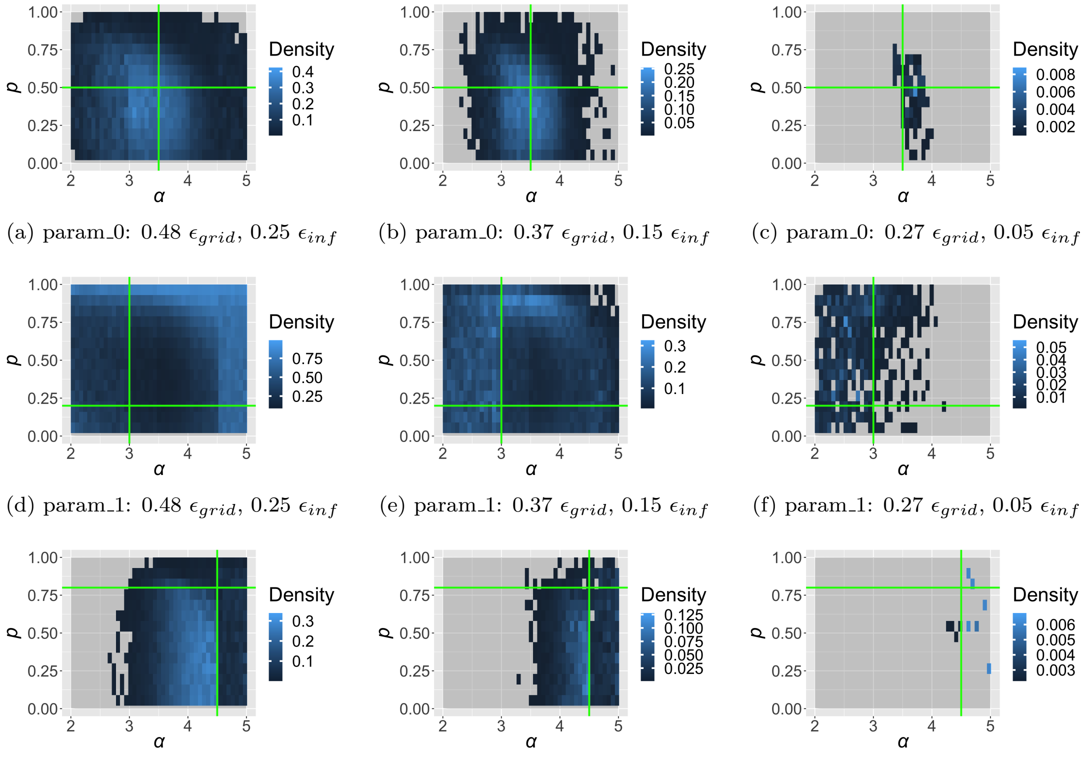


(a) param_0: 0.48 $\epsilon_{grid}$, 0.25 $\epsilon_{inf}$ (b) param_0: 0.37 $\epsilon_{grid}$, 0.15 $\epsilon_{inf}$ (c) param_0: 0.27 $\epsilon_{grid}$, 0.05 $\epsilon_{inf}$

(d) param_1: 0.48 $\epsilon_{grid}$, 0.25 $\epsilon_{inf}$ (e) param_1: 0.37 $\epsilon_{grid}$, 0.15 $\epsilon_{inf}$ (f) param_1: 0.27 $\epsilon_{grid}$, 0.05 $\epsilon_{inf}$

(g) param_2: 0.48 $\epsilon_{grid}$, 0.25 $\epsilon_{inf}$ (h) param_2: 0.37 $\epsilon_{grid}$, 0.15 $\epsilon_{inf}$ (i) param_2: 0.27 $\epsilon_{grid}$, 0.05 $\epsilon_{inf}$

Supplementary Figure 14: Recovering simulation parameters for three different input parameter sets using the three summary statistics in combination, $S_{cen}, S_{nat}, S_{grid}$, summarised as pairwise plots of $p$ and $\alpha$. The tolerances for both of the infectious proportion-based summary statistics, $\epsilon_{cen}, \epsilon_{nat}$, are represented by $\epsilon_{inf}$. (a-c) is parameter set 0, (d-f) is parameter set 1, (g-i) is parameter set 2.
